# Rule-based mitigation of charge asymmetry-triggered monoclonal antibody self-assembly

**DOI:** 10.1101/2025.11.03.686261

**Authors:** Inna Brakti, Anette Henriksen, Maria Łucja Tomczak, Stine Lyhne Godsk, Samuel Lenton, Vito Foderà, Minna Groenning

**Affiliations:** Department of Pharmacy, University of Copenhagen, Copenhagen, Denmark 2100; Biophysical Analysis, CMC Analytical Support, Novo Nordisk A/S, Måløv, Denmark 2760; Therapeutics Discovery, Novo Nordisk A/S, Novo Nordisk Park, Måløv, Denmark, 2760; API MSAT Stability, Novo Nordisk A/S, Bagsværd, Denmark 2880

**Author notes:** Vito Foderà, Address: Department of Pharmacy, University of Copenhagen, Copenhagen, Denmark 2100, Minna Groenning, Address: Biophysical Analysis, CMC Analytical Support, Novo Nordisk A/S, Måløv, Denmark 2760. **Author Contributions: I.B:** Conceptualization, Project administration, Investigation, Validation, Visualization, Writing - Original Draft, Formal analysis **A.H:** Resources, Investigation, Formal Analysis, Writing - Original Draft, Writing - Review & Editing, **M.L.T:** Investigation, Writing - Original Draft, **S.L.O:** Investigation, **S.L:** Project administration, Writing - Review & Editing, **V.F:** Conceptualization, Project administration, Resources, Supervision, Writing - Review & Editing, Funding acquisition, **M.G:** Conceptualization, Project Administration, Resources, Supervision, Writing - Review & Editing, Funding acquisition. **Competing Interest Statement**: A.H, M.L.T, S.L.G and M.G are employees of Novo Nordisk A/S.

**Keywords:** biopharmaceuticals, formulation, protein interactions, self-assembly, sub-visible particles, aggregation

## Abstract

In a pharmaceutical setting, understanding the factors governing monoclonal antibody (mAb) attractive interactions in formulations is highly warranted as many solution phenomena such as liquid-liquid phase separation (LLPS) result from their preferential self-interaction. While the effect of locally accumulated charge in the variable region has been recognized as an important factor in mediating non-specific mAb self-assembly, the effect of charge asymmetry, i.e. the distribution of oppositely charges residues, has been much less studied experimentally. Moreover, most studies restrict such analyses to the variable region of mAbs, leaving out possible contributions from the constant region of the molecule to the observed sticky behavior. Hence, the aim of this work is to correlate the charge asymmetry over the entire mAb surface to the extent of attractive self-interaction. To do so, we selected three mAbs with distinct solvent exposed surface distribution of charged residues, for which we computationally assessed the charge asymmetry and defined an apparent molecular stickiness ranking. We then tested this ranking experimentally by evaluating their ability to engage in attractive self-interaction as a function of mAb concentration and ionic strength. Experimental data included a combination of small-angle X-ray scattering, dynamic light scattering and micro-flow imaging. We show that the mAbs with oppositely charged Fab and Fc domains are characterized by overall attractive protein-protein interactions in solution amounting to diverse sub-visible morphologies, which vary non-linearly with mAb concentration and ionic strength. As a proof of concept, we also report the absence of any of such assemblies for the mAb with like-charged Fab and Fc domains, resulting in an overall repulsive behavior in solution. Altogether, we show how to utilize charge distribution analyses of full-length mAbs to rationally develop formulations that prevent problematic self-assembly.

**Graphical abstract:** 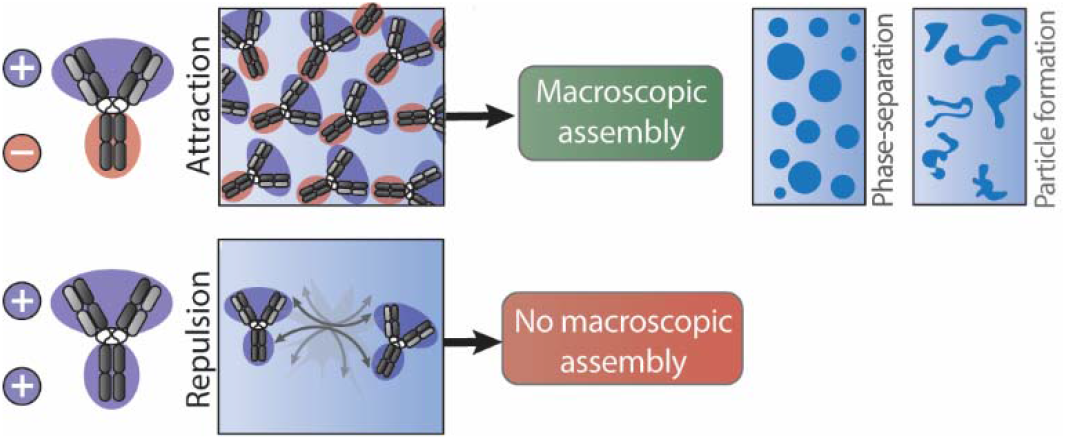

## Introduction

Therapeutic monoclonal antibodies (mAbs) represent the leading class of biopharmaceuticals with more than 200 marketed products for the targeted treatment of a wide range of diseases^1^. Moreover, the large number of mAbs and related modalities in development reflect the vast engineering possibilities this class of biotherapeutic molecules offers. In the context of chronic diseases, the delivery of mAb therapeutics via subcutaneous injection provides several patient-centric benefits including the convenience and flexibility of self-administration^2^. To accommodate the high doses and the small injection volume associated with this delivery route (maximum of ≈2 mL), therapeutic mAb concentrations usually exceed concentrations of 100 mg/mL^3^. The close molecular proximity and high propensity for intermolecular interactions associated with the very high concentrations can invoke a range of physical stability challenges such as high viscosity, opalescence, phase separation and aggregation^4^. The latter has been subject to investigation since some proteinaceous aggregates or particles on the sub-visible size range (2-100 µm) were shown to contribute to unwanted immunogenicity and severe adverse effects in humans^5–7^. In practice, identifying particles with immunogenic properties is complicated by the fact that distinct aggregate morphologies and internal structures can trigger different immune responses^8^. The complex interactions underlying mAb self-assembly remain poorly understood, complicating the development of predictive analyses and models that accurately capture their primary driving forces^9^. In a study from 2021, Lai et al. used logistic regression on a mAb viscosity dataset and found that the top three features in determining their viscosity profiles were the number of hydrophobic or hydrophilic residues in the variable (Fv) domains and the charge symmetric parameter of the entire mAb^10^, which is similar to the kappa (κ) value for disordered proteins^11^. Generally, they observed that the highly viscous mAbs in their dataset contained less hydrophobic residues and were enriched in hydrophilic ones. Indeed, the presence of local agglomerations of like-charged and polar residues in so-called patches along the mAb surface has been directly linked to developability issues resulting from reversible self-association of mAbs^12–17^. In support of this, mutational studies in which such surface patches were disrupted have on several occasions proven to be a successful strategy for abrogating problematic non-specific self-interaction^12,18–25^. As mentioned earlier, the colloidal stability of mAbs is also affected by their level of charge segregation as charge asymmetry along the mAb surface can lead to attractive self-complementary forces^13,14,26–29^. This is supported by several reports emphasizing the importance of both the magnitude and direction of the resulting dipole^14,17,30^. However, this property is usually reported for the variable domain and seldom studied for full-length mAbs^17^, despite experimental evidence showing that the presence of electrostatically complementary ‘sticker’ regions^31^ in the variable fraction and Fc domain can enhance their predisposition self-association^10,32–34^. The term ‘sticker’ was borrowed from associative polymer physics and introduced to the field of biomolecular phase transitions by Pappu and colleagues to describe molecularly adhesive regions, such as patches on folded proteins, whose synergy with the regions separating them (termed spacers), determine the overall self-association behavior^35^.

Several groups have brought forward a list of rules and guidelines for general therapeutic mAb developability screening, based on *in silico* analyses of large mAb datasets, which are reviewed elsewhere^15^. However, the preponderance of data available for IgG1 introduced a bias towards the inherent physicochemical properties of this particular isotype when deriving these rules, as noted by Agrawal et al.^36^. While representing a valuable starting point^15^, it is becoming increasingly clear that the different charge properties in the Fc domain across the IgG subtypes might complicate a direct translation of concepts learned from IgG1 studies to other isotypes. The constant region of lgG1 based mabs always bears a net positive charge caused by the high degree of positively charged residues present in the CH1 domains. A rational strategy to reduce self-interactions, thereby increasing solution behaviour, would therefore involve reducing the net negative charge and increasing the net positive of the variable region, resulting in an overall net positive charge of the whole mAb. This would increase the net repulsive interactions of the whole mAbs^10^. IgG4 mAbs, on the other hand have neutral C_H_1 domains and negatively charged Fc domains, most of which are concentrated in the Fc tail at formulation relevant conditions, as shown in the isopotential electrostatic surface generated by Skar-Gislinge et al.^37^. In this case, reducing positive charge in the variable region will reduce the size of the dipole along the mAb structure, and logically, the ability to self-interact via electrostatic complementarity^33^.

Here we study the effect of formulation conditions on intermolecular interactions and large-scale self-assembly of mAbs, based on their surface charge asymmetry properties. To achieve this, we selected three mabs representing distinct charge asymmetries and combined in-silico surface patch property analyses with small- and wide-angle X-ray scattering (SAXS and WAXS), dynamic light scattering (DLS) and micro-flow imaging (MFI) to study the effect of NaCl concentration and pH on intermolecular interactions and large-scale self-assembly. We show that the mAbs (mAb1 and mAb2) that have oppositely charged Fab and Fc regions can form higher order networks, while mAb3, with a more homogenous surface charge distribution, displays more repulsive behavior, resulting in an inability to self-assemble into larger species. Moreover, we highlight the fact that the protein-protein interactions (PPIs) leading to the formation of these assemblies at 45 mg/mL can be detected at a mAb concentration an order of magnitude lower. However, in this concentration regime, the different forces acting on the mAbs can result in assemblies with strikingly different morphologies, exemplified through mAb1 undergoing phase separation at 45 mg/mL, but forming sub-visible particles at 5 mg/mL. Finally, we provide a rational formulation strategy to minimize the extent of particle formation in low concentration mAb1 samples through modulation of the formulation pH, based on its anisotropic charge properties.

### Experimental section

#### Selected mAbs

The monoclonal antibodies, mAb1, mAb2 and mAb3 (two IgG4s and IgG1, respectively) were provided by Novo Nordisk A/S in proprietary formulations and stored as aliquots at -20°C which were thawed immediately before dialysis and only once, to avoid freeze-thaw cycles. All three mAbs have theoretically or experimentally determined isoelectric points of approximately 8.

#### Model generation of mAb1, mAb2 and mAb3

A structural model of a full antibody, mAb1, was generated by grafting available Fab crystal structures onto the corresponding IgG4 structures (PDB:5DK3). A C-termini capping of the crystal structure of mAb1 was applied in calculations of Fab properties for mAb1. C-termini capping was similarly applied to Fab homology models of mAb2 and mAb3 that were generated using the Antibody Modeler module in Molecular Operating Environment MOE (2024.06 release) ^38^ no crystals structures were available for these two molecules. The homology modelling was based on the default MOE antibody library applying default MOE settings for Fab homology modelling generating only one model for each molecule. The capping of the Fv C-termini was carried out to neutralize these and avoid their contributions to downstream calculations.

#### Fc domain net charge calculations

Net charge calculations for the Fc and Fc-hinge regions were carried out using the MOE protein property calculation module at pH 6.5, respectively MOE (2024.06 release) (26), and with the Fc structures extracted from the publicly available human IgG1 (PDB:1HZH) and human IgG4 (PDB:5DK3) structures. Default settings for the property calculations were used (dielectric constant set to 78, NaCl concentration set to 0.1 M).

#### Structure-based surface patch property calculations

We employed the methodology outlined by Ausserwöger et al.^21^ to compute the Fab surface patch property descriptors □_H_^Fab^, □_N_^Fab^ and □_P_. Specifically, the Protein Properties tool in MOE (2024.06) was utilized in ensemble mode generating 100 conformations of each model using the LowModeMD method^39^ without minimization to carry out constant-temperature conformational sampling of protonation states. Protonation states were assigned for the sampled conformations using the Protonate3D method^40^ with implicit Monte Carlo sampling adhering to the default settings for this tool. Samplings were centered at either pH 6.5 and 7.4 with NaCl concentration set to 0.10 M. An additional sampling was performed with 6.5 as pH midpoint but with the NaCl concentration adjusted to 0.01 M. As per default settings, hydrophobic patches consisted of regions where the hydrophobic potential was equal to or greater than that of a methyl group and persists over a surface area greater than 50 Å^2^. The hydrophobic potential was determined for each atom using the SLogP method and its result mapped onto the molecular surface. The charged patches were assigned where there was excess forcefield charge (Amber10:EHT) sustained over a surface area of at least 40 Å^2^. From this analysis, the negative surface patch area (□_N_) was then subtracted from the positive surface patch area (□_P_) to obtain the total charged patch area difference 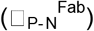.

#### Dialysis

A buffer solution consisting of 10 mM histidine was prepared using reagent grade L-histidine (ReagentPlus®, ≥99% TLC, H8000, Sigma Aldrich) in Milli-Q water (18 MΩ/cm at 25°C). The pH was then adjusted to 6.5 +/- 0.05 with HCl (Titripur®, reag. Ph. Eur., reag. USP, 1.09057,1000, Supelco) and the buffer was filtered using a 0.1 µm membrane (Stericup® Quick Release, Millipore Express® 0.10 μm PES, Cat no: S2VPU11RE). Buffer exchange into this buffer was carried out by dialysis, using 20K MWCO Slide-A-Lyzer G3 dialysis cassettes (Thermo Scientific™, Product number: A52976, Merck. The dialysis of each mAb was carried out separately with an exchange factor set to 500 for all steps of the dialysis. The mAb was dispensed into a humidified dialysis cassette and placed in dialysis for two hours at room temperature, with stirring. The buffer was then discarded and replenished before another dialysis for 2 hours at room temperature. Dialysis was repeated with overnight incubation at 4°C. All glassware used for the preparation of samples, and excipient stock solutions were thoroughly rinsed with three volumes of Milli-Q water and dried prior to use to minimize the introduction of foreign particulate matter.

#### Stocks and sample preparation

After dialysis mAb2, which was supplied at a lower concentration, was concentrated to 100 mg/mL using 10 kDa Amicon ultra centrifugal filters (Merck, Product number: UFC8010). From the three concentrated mAb stocks, dilutions were made in freshly prepared buffer (10 mM histidine pH 6.5). Fresh NaCl stocks were prepared at 40 mM (to prepare the 10 mM NaCl samples) and 500 mM (to prepare the 150 mM NaCl samples) due to the highly sensitive nature of mAb1 to NaCl concentration in the range 0-20 mM. Samples were always prepared by adding buffer first, followed by NaCl, then the mAb stock and immediately thereafter inverting the microcentrifuge tube. All samples were filtered before analysis using a 0.1 µm Whatman Anotop filter (GE Healthcare, product number: 6809-2012) connected to a silicone-free Luer lock syringe (Henke Sass Wolf, Henke Ject®, product number: 4050-457 X00V0) pre-washed as described in^41^.

#### Sample concentration determination

The absorbance at 280 nm was determined for a mAb dilution series using a NanoDrop™ One C spectrophotometer to avoid non-linear concentration effects. Absorbance values for dilution factors in the linear region of the mAb concentration versus dilution factor plot were multiplied by the dilution factor and the concentration in mg/mL determined using the theoretical extinction coefficients and molecular weights of the mAb before being averaged.

#### SAXS and WAXS

Small and Wide-angle X-ray scattering (SAXS and WAXS) experiments on 45 mg/mL mAb1 and mAb3 in 10 mM histidine + 0/10/150 mM NaCl samples at pH 6.5 were conducted at room temperature at the CoSAXS beamline at the MaxIV laboratory (Lund University). The capillary was initially washed in a 10 mM histidine pH 6.5, 150 mM L-arginine HCl buffer, followed by a wash with matched sample buffer before measuring a freshly prepared, matched buffer for background subtraction. The scattering intensity of the mAb samples were measured immediately after. An additional buffer measurement was performed post-sample to ensure that no protein buildup occurred in the capillary. For all buffer and sample measurements, forty consecutive frames of 50 ms were collected and placed on absolute scale using water and empty capillary measurements. Prior to averaging, consecutive frames were investigated for radiation damage, after which the corresponding background was subtracted. The scattered beam was detected on a Eiger2 4M detector for SAXS and a Pilatus L-shaped 2M Mythen 12 detector for WAXS after which the isotropic scattering was azimuthally averaged to produce the one-dimensional scattering curve of the intensity (a.u) versus the scattering vector q= 4*sin(θ)/λ, where θ is the half scattering angle and λ, the wavelength of the X-rays. All subsequent data analysis was performed using the Primus package available in the ATSAS distribution^42^.

#### Dynamic light scattering

The apparent hydrodynamic radius was measured as independent duplicates at room temperature (22 °C), using dynamic light scattering (DynaPro® III Plate Reader, Waters, Wyatt Technology, Santa Barbara, USA). A sample volume of 25 µL was pipetted into 384 well Aurora Microplates (Whitefish, USA). The microplate was sealed with tape (ThermoFisher, Product number; 235307) and centrifuged for 5 minutes at 1780 *g*. The acquisition time was set to 5 sec and the acquisitions per sample to 40 as described in^43^. The autocorrelation function was fit by cumulant analysis to extract the hydrodynamic radius.

#### Sub-visible particle analysis using Micro-Flow Imaging

Flow imaging analysis using Micro-Flow Imaging™ was used for the quantification of sub-visible particles and analysis of their morphologies as a function of formulation conditions. Samples were prepared and analyzed in 96 deepwell plates (Waters, cat. number 186002482) on a Micro-flow imaging™ 5200 particle analysis instrument equipped with an autosampler (ProteinSimple/Bio-Techne, San Jose, California, USA). First, two sets of two system flushes were carried out at maximum speed with Milli-Q water. Next, the uncoated flow cell with depth of field-matched flow cell depth of 100 μm (part number 4002-005-001, ProteinSimple, Toronto, ON, Canada) was cleaned with 1% Tergazyme/0.5% Liquinox (v/v) filtered through a 0.1 µm-filter (Stericup Quick Release, Millipore Express, cat. number S2VPU11RE, Merck) and Milli-Q water until the recorded particle concentration was below 150 particles/mL. At this threshold, the instrument was considered clean. Subsequently, the sample was resuspended before aspiration and introduction into the sample port for analysis. The initial 220 μL of the sample were utilized to optimize the illumination settings of the camera, thereby enhancing particle detection within the designated field of view. Following this, approximately 150 µL of the sample was employed for system purging, while around 260 µL underwent analysis at a flow rate of 0.17 mL/min. Between individual sample analyses, the flow cell was subjected to a cleaning protocol involving two flushes with milli-Q water, followed by two flushes with a cleaning solution comprising 1% Tergazyme and 0.5% Liquinox (v/v), and concluded with an additional two flushes utilizing milli-Q water. Data processing was carried out automatically using in-house software which ensures the removal of slow/stuck (and consequently, imaged multiple times) particles combined with an algorithm designed to improve the detection of translucent particles, if needed^44^. The data was inspected to make sure that the volume of sample used for flow-cell purging was sufficient to avoid any dilution effects from the cleaning solutions. Additional filters were applied in the analysis software to remove and only consider particles in the 5-1000 um range. The data quality of the final dataset was evaluated based spatial randomness statistics provided by the software. Scatter plots were further examined to detect any inhomogeneities that might interfere with the analysis.

## Results

### mAb self-assembly is dependent on complementary sticker regions

We first modelled the surface patch distribution in the Fab domains and calculated the difference between positive 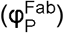 and negative 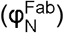 charged patches to get an idea of the overall charged patch area difference 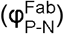, as performed in a previous study ^21^. This analysis established that all three mAbs contain an overall excess positive charge in their Fab domains, the size of which decreased from mAb1 to mAb3 (Figure 1). Additionally, all three mAb Fab domains were characterized by the presence of a hydrophobic patch of comparable size.

**Figure 1:**
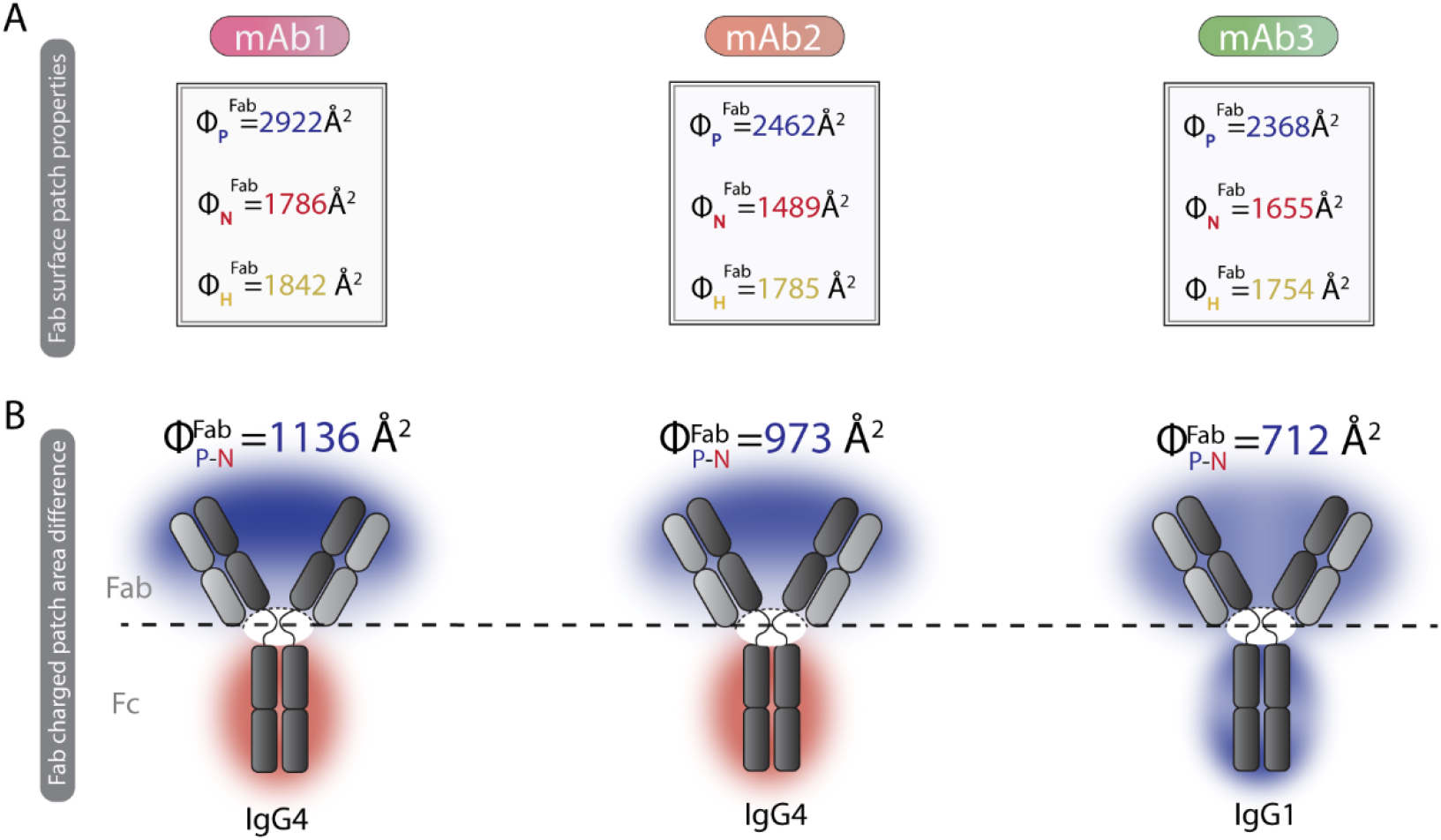
Surface patch properties of mAb1-3 in 10 mM histidine pH 6.5 + 10 mM NaCl. (A) The total positive, negative and hydrophobic surface patch area (□_P,_ □_N_ and □_H_, respectively) for the entire Fab region determined using MOE (2024.06 release) are reported in Å^2^ for all three mAbs in the top panel. (B) Total charged patch (□_P-N_^Fab^) area difference in the Fab region of the different mAbs, are emphasized with a blue shading on the Fab domains as a result of their overall positive charge. The hinge region, highlighted by the dashed line on the illustration of the mAbs, and the Fc domains were excluded from this part of the analysis. The Fc domains are shaded according to their calculated net charge properties (negative: red, positive: blue). See values in Table S1.

To compare the charge properties in the Fab domain with the rest of the molecule we also calculated the net charge properties of the IgG4 and IgG1 Fc domains at pH 6.5 (Table S1). Both mAb1 and mAb2 are of the IgG4 subtype and have a negatively charged Fc domain at pH 6.5, in which the negative charge patch in the Fc tail represents an obvious complementary sticker region for the positive surface patch in the Fab domain^33,37^. As described for other mAbs, segregation of charges along the mAb surface is instrumental in granting them the ability to engage in homotypic Fab-Fc interactions^33,37,45^. On the other hand, mAb3, an IgG1 mAb, has a positive Fc domain and is therefore expected to display a higher degree of electrostatic repulsion and a lower tendency for self-assembly, compared to mAb1 and mAb2. From our previous study^46^, mAb1 was shown to have an intricate response to ionic strength, undergoing liquid-liquid phase separation (LLPS) in a very narrow NaCl concentration range at 45 mg/mL of mAb1, while a slight increase in the ionic strength preserved the highly opalescent appearance of the solution but abolished the formation of droplets. When supplied with higher NaCl concentrations the mAb1 solution transitioned back to a transparent, well mixed solution, which is a typical feature of proteins with oppositely charged surface patches^14,16,29,37^.

As a first step, we tested the validity of our hypothesis that charge-distribution is accompanied by the presence of attractive/repulsive interactions by first studying mAb1 and mAb3, which represent the two extreme cases. Based on our charge asymmetry analysis (Figure 2) we hypothesize that mAb1 and mAb3 should show opposite behaviours. To this end, we acquired high-resolution SAXS and WAXS data which report on both colloidal and conformation stability, without the need of prior sample dilution^9,37,47,48^.

**Figure 2:**
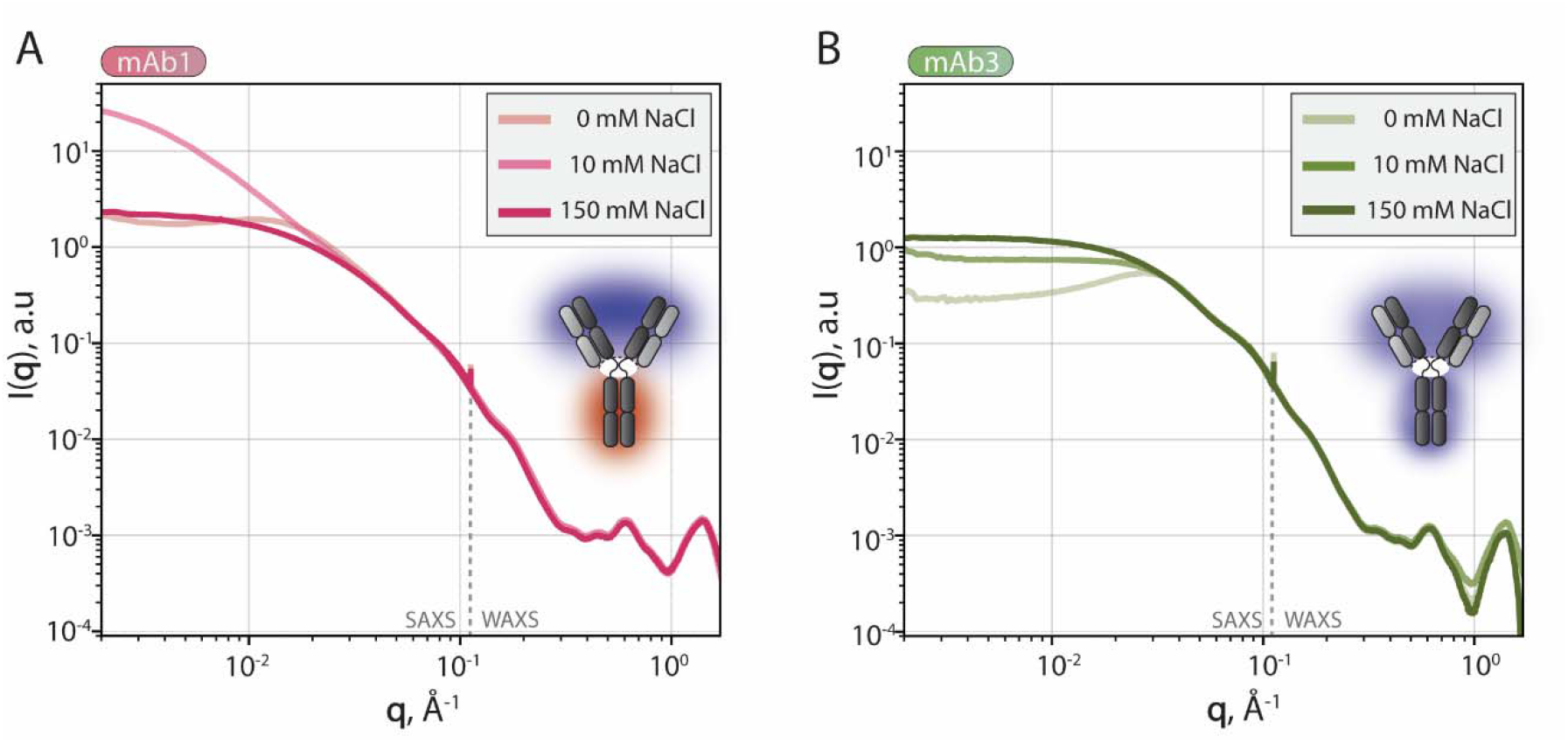
SAXS and WAXS profiles for mAb1 and mAb3 as a function of NaCl concentration. Scattering curves (intensity as function of scattering vector, **q**) of 45 mg/mL (A) mAb1 and (B) mAb3 samples in 10 mM histidine pH 6.5 containing 0, 10 or 150 mM NaCl (increasing color intensity).

This aspect is particularly important for mAbs as they are typically formulated at high concentrations where the macroscopic repercussions of non-specific self-assembly such as high viscosity, opalescence and phase transitions usually manifest^9,26,37,49^. In a SAXS curve, structural features as well as interparticle interactions can be probed, the former producing noticeable distortions in the low-q region of the plot of I(q) vs q^50^.

As shown in Figure 2, we observed pronounced effects of NaCl concentration on the scattering intensity at low q of mAb1, as was shown in our previous study^46^. At 0 mM NaCl the scattering curve displayed an unusual shape, with a visible downturn at q ≈ 0.004 Å, resulting in an upturn of the signal at even lower q-values. At 10 mM NaCl, a sharp increase in the scattering intensity was observed, resulting from the formation of micron-sized, concentrated droplets, while at 150 mM the signal decreased substantially although a slight structure factor contribution could be discerned. For mAb3 on the other hand, a downturn in the intensity at low angles could be seen for the 0 mM NaCl condition, indicating that interactions are mainly repulsive under these ionic strength conditions. As the salt concentration was increased to 150 mM, the scattering intensity at low q increased until roughly intersecting the y-axis perpendicularly, suggesting that the extent of PPIs was minimal. For both mAbs, the scattering curves showed a high degree of overlap at higher q values, indicating that the conformation of the mAb was likely unaltered by the addition of NaCl. Altogether, these results are in line with our predicted self-interaction propensity from the surface charge properties (e.g., mAb1 and mAb3 showing attractive and repulsive interactions, respectively) albeit with different responses to NaCl concentration.

Next, we studied the dependence of the PPIs as a function of the mAb and NaCl concentration by monitoring the DLS signal. We specifically focused on a mAb1/2/3 dilution series 45 to ≈1.4 mg/mL at 0, 10 or 150 mM NaCl (Figure 3). In this setup, we included mAb2, which, according to our surface patch analysis, has a less positive Fab compared to mAb1 and a negative IgG4 Fc tail^37^. Based on this stickiness ranking, mAb2 is expected to display an intermediary self-association behavior. Here ⟨*R*_*h*_⟩_,*app*_ refers to the apparent population averaged hydrodynamic radius derived from the Stokes-Einstein equation D = k_B_T/6πηR_h_, where D is the ensemble diffusion coefficient, k_B_ is the Boltzmann constant, T is the absolute temperature, and η is the solution viscosity (see Tables S2-S4 for exact values). As shown in Figure 3, the three mAbs had different responses to the NaCl concentration. At 0 mM NaCl, the ⟨*R*_*h*_⟩_,*app*_ of mAb1 slightly increased from the expected size of an individual antibody (≈5-6 nm)^43^ to 7.2 nm at 45 mg/mL. The opposite was observed for mAb3, whose apparent size decreased from roughly 4.6 to 2.3 nm. This artificially low molecular size measured at higher concentrations is consistent with an increased repulsion within the system and aligns with the noticeable downturn in scattering intensity at low scattering angles observed in Figure 2^51^. When increasing the NaCl concentration from 0 to 10 mM, the ⟨*R*_*h*_⟩_,*app*_ of mAb1 underwent a drastic increase, reaching approximately 55 nm at 45 mg/mL. This was expected as this mAb was shown to undergo phase separation above 30 mg/mL under these conditions^46^. At 10 mM NaCl, the size of mAb3 was roughly that of a monomeric mAb and remained virtually unchanged as a function of mAb concentration, decreasing only from 5 to 4.6 nm (see Table S4), suggesting that some of the repulsion at 0 mM NaCl had been suppressed and balanced by an attractive interaction potential^51^.

**Figure 3:**
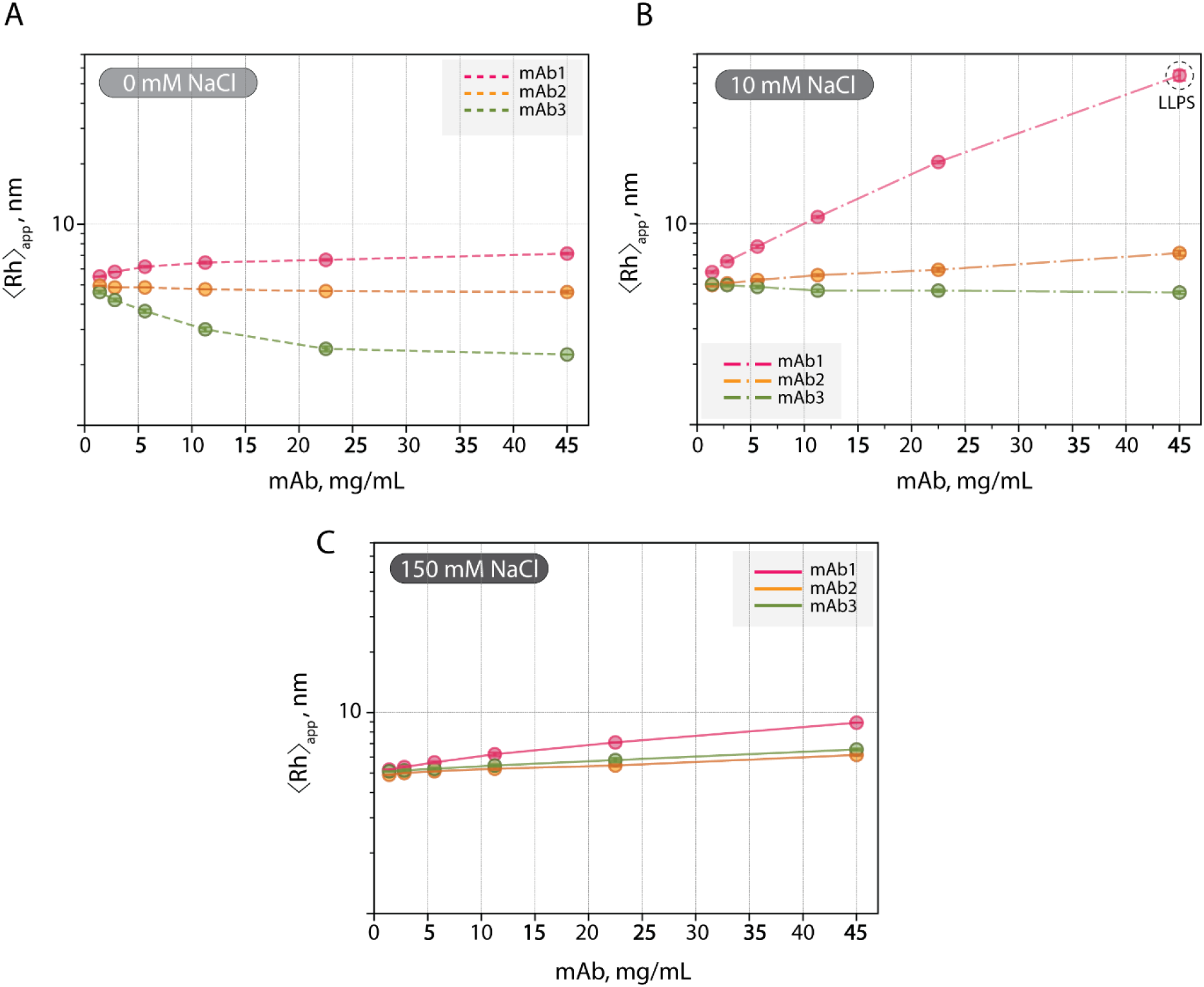
Dynamic properties of mAb samples as a function of NaCl and mAb concentrations. Apparent hydrodynamic radii (⟨*R*_*h*_⟩_,*app*_) of (A) mAb1 (pink), (B) mAb2 (orange) and (C) mAb3 (green) dilution series as a function of mAb concentration (≈1.4 to 45 mg/mL) and NaCl concentration (0,10,150 mM), measured using DLS. Error bars represent standard deviations from duplicate measurements but are in most cases not visible as they are too small.

In most cases, mAb2 displayed a behavior positioned between the responses observed for mAb1 and mAb3 (see Table S3). At 0 mM NaCl, the ⟨*R*_*h*_⟩_,*app*_ underwent a modest decrease in size as a function of mAb concentration (5 to 4.6 nm), indicating a weak repulsion within the system. By adding 10 mM NaCl to the solution, these interactions then switched from slightly repulsive to slightly attractive as inferred from the small increase in ⟨*R*_*h*_⟩_,*app*_ from 5 to 7.2 nm. Lastly, at the highest NaCl concentration of 150 mM, all three mAbs showed an increase in ⟨*R*_*h*_⟩_,*app*_, albeit to different extents. mAb1 showed the largest effect, reaching an apparent size of 8.9 nm at 45 mg/mL, while a less pronounced increase in the hydrodynamic radius was observed for mAb2 (4.9 to 6.2) and mAb3 (5.1 to 6.6). Interestingly, the observed trends were already detected at low mAb concentrations, prompting us to evaluate whether the increase of ⟨*R*_*h*_⟩_,*app*_ was solely due to attractive interactions as observed by Sukumar et al.^52^ or if they also translated into the formation of detectable mAb assemblies.

Here, we used micro-flow imaging (MFI) to detect the presence of such large-scale assemblies in the subvisible size range from 2-100 µm. In this setup, an illuminated sample flows through a thin flow cell positioned in the field of view of a microscope, during which images are continuously acquired^53^. Particles with refractive indices differing from that of the solution appear with a reduced light intensity, allowing their automated detection and morphology analyses as well as quantitative analyses^44,53,54^. Enhanced detection of translucent particles, which are commonly observed for proteins, can further be achieved using more advanced algorithms^44^.

As presented in Figure 4A (and Figures S1 and S3), mAb1 samples at 45 mg/mL were characterized by a few small particles at 0 mM and 150 mM, with larger and more translucent particles being formed in the latter, which explains the slight structure factor contribution in the low-q region of the SAXS curve and higher hydrodynamic dimensions recorded for the sample at high salt concentration (Figures 2A and 3C). At 10 mM NaCl, spherical droplets were detected in the sample, consistent with the phase separation observed using orthogonal techniques^46^ (Figure S2). Strikingly, at 5 mg/mL, large particles were observed at all three ionic strength conditions, with translucent worm-like morphology being formed almost exclusively at 0 mM NaCl (Figures 4A, S4) while darker, more compact morphologies occurred at 10 and 150 mM NaCl (Figures 4A, S5-6). Similarly, mAb2 formed large assemblies close to the visible range at 45 mg/mL, but with a difference NaCl-dependency compared to mAb1. Indeed, larger particles were observed at 0 and 150 mM NaCl compared to 10 mM NaCl, highlighting the non-linear nature of particle formation with respect to the NaCl concentration (Figures 4B, S7-10). At 5 mg/mL, large translucent assemblies were detected for all three NaCl conditions (Figures 4B, S11-13). For mAb3, the mesoscale self-assembly profile, assessed using MFI, was invariant throughout the NaCl and mAb concentrations tested, with only small particles detected (Figures 4C, S14-15).

**Figure 4:**
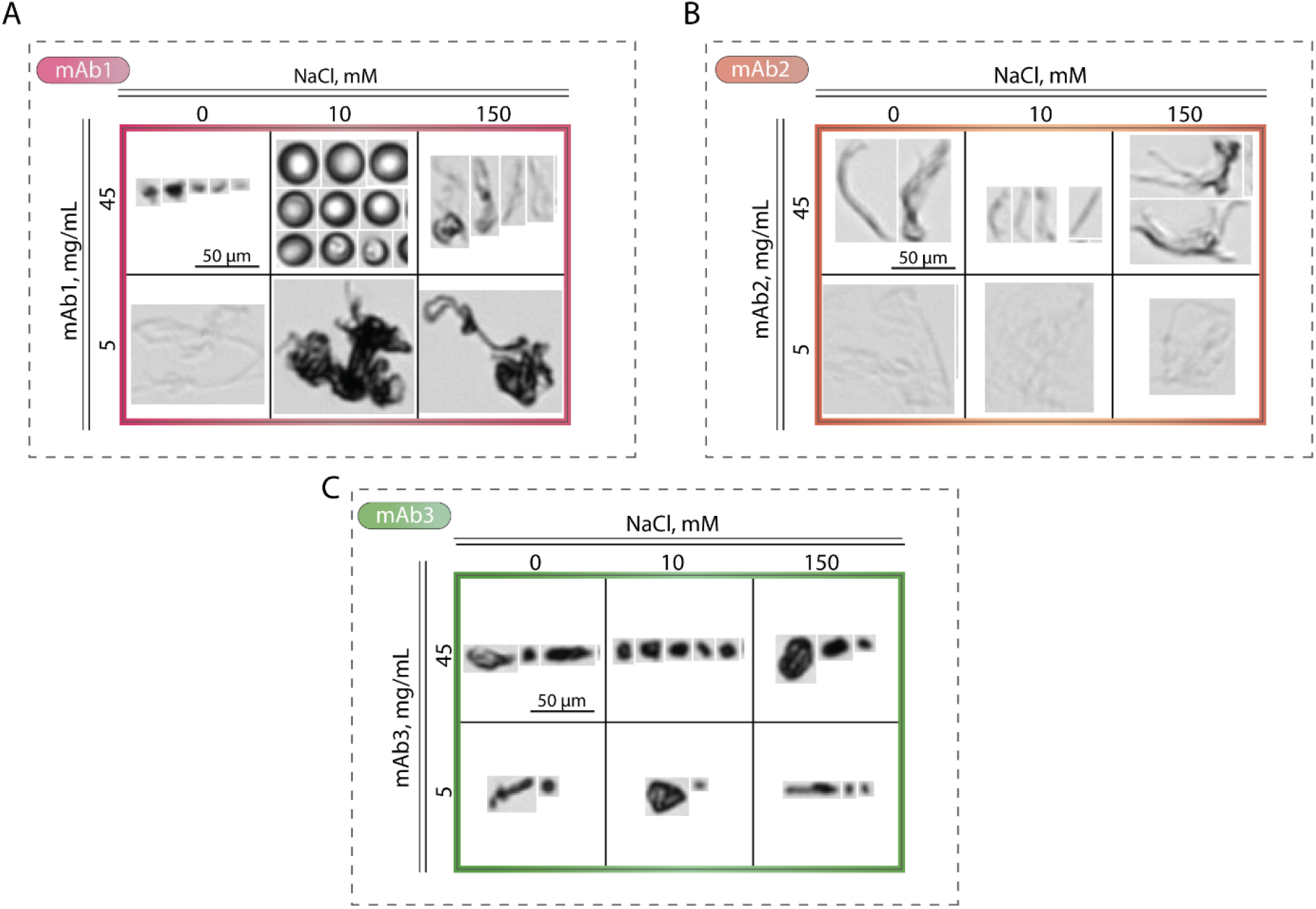
Evaluation of particle morphology using micro-flow imaging. (A) mAb1 (pink), (B) mAb2 (orange) and (C) mAb3 (green) samples at 45 and 5 mg/mL in 10 mM histidine pH 6.5 were evaluated for their ability to form large-scale assemblies as a function of NaCl concentration (0/10/150 mM). Representative images of the largest particles formed are reported here for ranking purposes. A scale bar is included in the first image, which applies to all images.

### Surface properties as guides for rational formulation design

During drug development, simple formulations containing as few excipients as possible, while preserving the integrity of the active pharmaceutical ingredient are highly warranted. In this final part, we use the example of mAb to show that the extent of attractive interactions in solution can be achieved without the addition of excipients, by simply altering the solution pH. Based on the charge properties of mAb1, we hypothesized that lowering the solution pH to 5 would neutralize the side chains of acidic residues in the Fc domain and thereby decrease the dipole along its structure. To evaluate the effect of decreasing the pH from pH 6.5 to 5 we again turned to MFI and assessed particle morphology at 45 and 5 mg/mL mAb1, as a function of NaCl concentration (0,10 and 150 mM). As shown in Figure 5 (and Figures S16-17), shifting the solution pH to 5 completely inhibited the LLPS observed at 45 mg/mL for mAb1, which has been linked to the aberrant aggregation of some protein systems^55,56^. Moreover, despite a few particles being detected at 5 mg/mL mAb, none reached the large sizes observed at pH 6.5 sample supporting that pH modulation can induce morphological changes in mAb formulations and that it be an efficient strategy to minimize self-association propensity without the addition of excipients.

**Figure 5:**
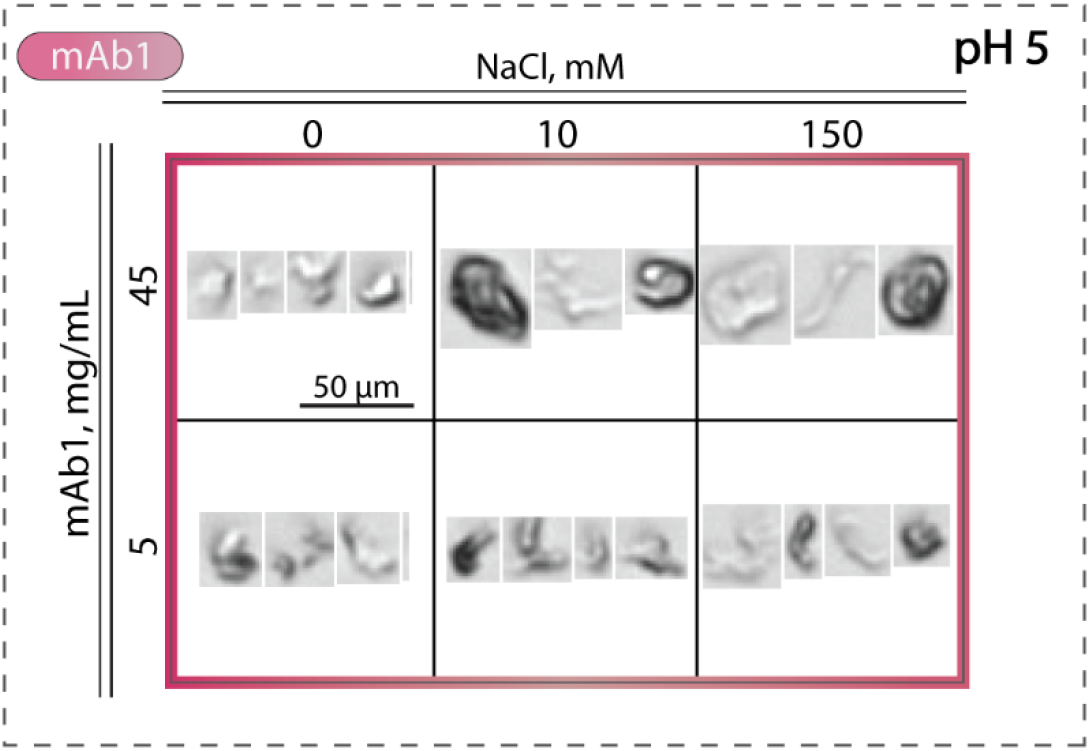
Evaluation of mAb1 particle morphology at pH 5 using MFI. Images of the largest particles formed are reported here for ranking purposes. A scale bar is included in the first box, which applies to all images.

## Discussion

### Surface charge properties as coarse predictors of self-assembly propensity

In this study, we aimed to increase our understanding of the molecular determinants of mAb self-assembly by performing a head-to-head comparison of three different mAbs with varying charge polarity along their total surface. Since the formulation conditions are identical for the mAbs, the difference in the stability behavior should uniquely arise from the intrinsic properties of the molecules. We hypothesized that a potential reason for the observed difference was related to the distribution of charges on the molecules. More specifically, we speculated that the root cause of this behavior was based on self-complementing electrostatic forces between the Fab and Fc domains and ranked the three mAbs according to their self-assembly propensity based on a surface charge distribution analysis (Figure 1). It is worth mentioning that we restricted the surface patch evaluation to the Fab domain and computed the net charge of the IgG1 and IgG4 Fc domains and referred to published potential surfaces for the distribution of the charges herein^37,51^. Indeed, we found that the two mAbs with self-complementary Fab and Fc regions (mAb1 and mAb2) could undergo large-scale self-assembly, which was orchestrated by NaCl in a non-linear manner, with mAb1 (Figure 4). Conversely, mAb3 displayed more repulsive interactions at 0 mM NaCl (Figures 2 and 3), becoming slightly more attractive as the NaCl concentration reached 150 mM, but not to an extent conducive to the formation of assemblies such as the phase separated droplets or large sub-visible particles observed for mAbs 1 and 2 (Figure 4). We hypothesize that the dominating forces driving the interaction landscape of the mAbs are related to charge asymmetry along the mAb surface and the generation of a dipole, the magnitude of which further increases the stickiness of the molecule and self-assembly propensity. However, we did not account for the distribution of such patches but envision that their specific location and accessibility can drastically influence the ability to engage in intermolecular interactions. Moreover, hydrophobicity has on several occasions been predicted and experimentally demonstrated to play a key role in the self-assembly of mAbs^10,21^. In this study, all three mAbs have a comparable total hydrophobic surface and hydrophobic patches were therefore not explicitly accounted for. However, the distribution of these patches on the mAb surface could vary between them and, depending on their location, they could act as complementary sticky regions. In Figure 3D, we summarized the effect of NaCl concentration in the evolution of the apparent hydrodynamic radius of the mAbs at a fixed concentration of 45 mg/mL. Both mAb1 and mAb2 experienced an increase in apparent ⟨R_h_⟩_,app_ at 10 mM NaCl, before decreasing at 150 mM. The observed effect was however much more pronounced for mAb1 than mAb2. A similar behavior has been reported for a range of proteins with anisotropic charge distributions, resulting in their preferential alignment and self-assembly via these complementary charged patches^29,33,57,58^. In such systems, the slight screening provided by a small increase in ionic strength reduces the Debye Length and weakens the Coulomb repulsion resulting from the net charge of the proteins^59^. This increases the effect of the dipole induced by the patches of opposite charges and, by extension, the self-interaction propensity even in the presence of a net charge^24^. By further screening the electrostatic interactions with higher salt concentrations, we hypothesize that these attractive charged-patch interactions are gradually suppressed, leaving space for other contributions, e.g. hydrophobic interactions. For mAb3, the increase in attractive interactions at 150 mM NaCl could therefore possibly be explained by a favorable interaction mediated by e.g. the hydrophobic patch in the Fab domain (Figure 1). Additional factors that may influence the range of accessible PPIs of the various mAbs include the inherent structural differences between the IgG1 and IgG4 constant regions, which were not addressed in our analysis. However, differing dynamics caused by e.g. different disulfide bond patterns^60^ and/or the disordered hinge region, which varies in length, sequence, charge properties and flexibility between IgG1 and IgG4, could further modulate the interaction potentia^61^. Interestingly, IgG4 Fab domains have been shown to contact the C_H_2 domain of the Fc via the C_H_1 domain. This ability could place complementary sticky regions in contact and promote or shield self-assembly in a concentration-dependent way^62^. Moreover, therapeutic mAbs are routinely optimized through engineering, which introduces additional structural aspects that can affect their stability. As an example, the IgG4 hinge usually contains a single-point mutation to mitigate its tendency to form half-antibodies and bispecific antibodies by Fab arm exchange^62^. Finally, in some cases, specific amino acids have been shown to play a non-negligible role in mAb self-interaction^10,29,63^, as well as direct binding of solvent components, such as ions^64^ and buffering agents^65,66^.

As previously highlighted, the developability of mAbs can be challenged by high viscosity, LLPS and aggregation. Aggregation of therapeutic proteins and particle formation have in some cases been linked to increased immunogenicity. These phenomena can occur during manufacturing, storage, transport and patient administration and can be induced by external stress, but also because of formulation parameters such as ionic strength, pH and temperature^43,44,67^. Therefore, identification of the root cause of particle formation as well as the development of strategies to minimize their occurrence are warranted. Generally, the macroscopic manifestations of mAb self-assembly are witnessed at higher protein concentrations where the intermolecular distances are reduced and interaction potentials enhanced. Accordingly, we observed that mAb1 and mAb2 formed large-scale assemblies with different morphologies (LLPS and/or particles) at 45 mg/mL (Figure 4A and B). The attractive interactions driving the transition into these assemblies were evident at much lower concentrations. This observation led us to investigate if the decreased proximity of the mAbs in the 5 mg/mL regime and the resulting physicochemical properties of the solution could lead to the formation of assemblies with different morphologies. Indeed, far below the critical saturation concentration for mAb1 LLPS, at 5 mg/mL, large dark particles were observed (Figure 4A). This suggests, that despite the nature of the interactions being the same in low and high concentration regimes, then the resultant mAb assemblies differs. Interestingly, at 45 mg/mL, a mAb1 solution without any added salt contained virtually no particles. However, at 5 mg/mL, large translucent particles were observed, highlighting that self-association and aggregation are highly non-linear and that lowering the mAb concentration might not always produce the desired effect. Interestingly, despite the large size of the particles formed at the latter condition, only modest effects on the hydrodynamic radius were observed, which could be explained by concentration and possible sedimentation of the particles (Figure 3). However, a quantitative assessment of the particles formed was not the scope of the present work.

In addition to providing useful guidelines for the selection of well-behaved candidates in the early discovery process, understanding the effect of molecular surface properties on self-assembly can also guide formulation optimization campaigns at later stages. As pinpointed by Thorsteinson et. al.^30^, circumventing undesirable biophysical properties of lead candidates with desirable biological function can be achieved by altering key formulation parameters such as the pH^15^. For sticky IgG4 molecules, such as mAb1 and to a lesser extent mAb2, capable of undergoing large-scale assembly, lowering the solution pH represents a convenient way of neutralizing their negative Fc domain and thereby disrupting the dipole along the mAb surface. This effectively inhibited the phase-separation of mAb1 at 45 mg/mL and considerably reduced the sizes of the particles observed in solution, highlighting the possible benefits of designing formulations tailored to the needs of individual mAbs based on their molecular properties (Figures 5, S16-17). However, while low pH values can improve physical stability, the more acidic environment can enhance the rate of chemical degradation reactions such as deamidation, which can reduce binding affinity if located in the complementarity determining region of the mAbs^68,69^. Moreover, chemical modifications can also be accompanied by structural changes and hence affect physical stability. As such, formulation development is ultimately a compromise between physical and chemical stability.

## Conclusion

In this study, we show the agglomeration of like-charges into patches along the mAb surface steer their self-assembly behavior, despite a net charge on the molecule. Consequently, in addition to the amount of charged residues, considering their distribution over the total mAb surface is important to more accurately predict general mAb self-assembly as a function of formulation conditions. We further show that attractive interactions can result in different morphologies depending on the mAb concentration and ionic strength and more specifically that this relationship is highly non-linear. This entails that the behaviour observed at low concentration might not always extrapolate to higher concentrations and vice versa, which should be taken into consideration when studying mAb physical stability^70^. Despite mAb physical stability being complex and multifactorial, we hypothesize that simple *in silico* assessments such as the one performed here can help flag possible unwanted self-assembly early in the development process. This will avoid resource-demanding root-cause analyses of suboptimal physical stability profiles witnessed at later stages in the development pipeline.

## Abbreviations

DLS: Dynamic Light Scattering
Fab: Fraction Antigen Binding
Fc: Fraction Crystallizable
Fv: Fraction Variable
LLPS: Liquid-Liquid Phase separation
mAb: Monoclonal Antibody
MFI: Micro-flow Imaging
PPI: Protein-Protein Interaction
⟨*R*_*h*_⟩_,*app*_: Apparent Hydrodynamic Radius
SAXS: Small-angle X-ray Scattering
WAXS: Wide-angle X-ray Scattering

## Supporting Information

- Table S1: Calculated net charge of IgG1 and IgG4 Fc domains
- Table S2-S4: DLS data of mAb1-3 dilution series as a function of NaCl concentration.
- Figures S1-S3: MFI data of 45 mg/mL mAb1 solutions containing 0,10 or 150 mM NaCl at pH 6.5
- Figures S4-S6: MFI data of 5 mg/mL mAb1 solutions containing 0,10 or 150 mM NaCl at pH 6.5
- Figures S7-S10: MFI data of 45 mg/mL mAb2 solutions containing 0,10 or 150 mM NaCl at pH 6.5
- Figures S11-S13: MFI data of 5 mg/mL mAb2 solutions containing 0,10 or 150 mM NaCl at pH 6.5
- Figure S14: MFI data of 45 mg/mL mAb3 solutions containing 0,10 or 150 mM NaCl at pH 6.5
- Figure S15: MFI data of 5 mg/mL mAb3 solutions containing 0,10 or 150 mM NaCl at pH 6.5
- Figure S16: MFI data of 45 mg/mL mAb1 solutions containing 0,10 or 150 mM NaCl at pH 5
- Figure S17: MFI data of 5 mg/mL mAb1 solutions containing 0,10 or 150 mM NaCl at pH 5

## Acknowledgements

Novo Nordisk A/S is thanked for the funding this project as well as for providing the mAb material used to carry out this study. Jesper Søndergaard Marino from Novo Nordisk A/S is thanked for providing the MFI image evaluation software. VF, SL and IB acknowledge the support from the VILLUM FONDEN by the Villum Young Investigator Grants “Protein superstructure as Smart Biomaterials (ProSmart)” 2018-2023 (project number: 19175) and “Protein Phase Separation and Solid Transition in Synthetic Cells (ProSeC) 2024-2027 (project number: 53132). VF and SL acknowledge the Novo Nordisk Foundation (projects NNF20OC0065260 and NNF22OC0080141) for financial support.

