## Supplementary data for "Rule-based mitigation of charge asymmetry-triggered monoclonal antibody self-assembly"

* Vito Foderà

Address**:** Department of Pharmacy, University of Copenhagen, Copenhagen, Denmark 2100

Phone:

****

* Minna Groenning

Address: Biophysical Analysis, CMC Analytical Support, Novo Nordisk A/S, Måløv, Denmark 2760

****

| Format | Net charge pH 6.5 |
| --- | --- |
| Human IgG1 Fc (PDB:1HZH) | +9.1 |
| Human IgG4 Fc (PDB:5DK3) | -1.8 |

**Table S1: Net charge properties of the human IgG1 and IgG4 Fc domains calculated using MOE protein property calculation module at pH 6.5, respectively MOE (2024.06 release)**^38^

| **[NaCl]** | **DF**  **(mAb1)** | ***〈R_h_〉_,app_* _replicate_1** | ***〈R_h_〉_,app_* _replicate_2** | **Average** | **Std** |
| --- | --- | --- | --- | --- | --- |
| **0** | 1 (45 mg/mL) | 7,2 | 7,1 | 7,2 | 0,07 |
| **0** | 2 | 6,7 | 6,6 | 6,7 | 0,07 |
| **0** | 4 | 6,5 | 6,4 | 6,5 | 0,07 |
| **0** | 8 | 6,2 | 6,1 | 6,2 | 0,07 |
| **0** | 16 | 5,8 | 5,8 | 5,8 | 0,00 |
| **0** | 32 | 5,5 | 5,5 | 5,5 | 0,00 |
| **10** | 1 (45 mg/mL) | 57 | 53 | 55 | 3,2 |
| **10** | 2 | 21 | 20 | 20 | 0,28 |
| **10** | 4 | 11 | 11 | 11 | 0,14 |
| **10** | 8 | 7,8 | 7,6 | 7,7 | 0,14 |
| **10** | 16 | 6,5 | 6,4 | 6,5 | 0,07 |
| **10** | 32 | 5,8 | 5,7 | 5,8 | 0,07 |
| **150** | 1 (45 mg/mL) | 8,9 | 8,9 | 8,9 | 0,00 |
| **150** | 2 | 7,1 | 7,1 | 7,1 | 0,00 |
| **150** | 4 | 6,3 | 6,1 | 6,2 | 0,14 |
| **150** | 8 | 5,7 | 5,6 | 5,7 | 0,07 |
| **150** | 16 | 5,4 | 5,3 | 5,4 | 0,07 |
| **150** | 32 | 5,2 | 5,2 | 5,2 | 0,00 |

**Table S2: Apparent hydrodynamic radii (*〈R_h_〉_,app_*) of a mAb1 dilution series (dilution factor (DF) 1 (corresponding to 45 mg/mL) to 32, as a function of NaCl concentration ([NaCl].** The average was determined from two independent samples (*〈R_h_〉_,app_* _replicate_1 and *〈R_h_〉_,app_* _replicate_2), from which the

standard deviation (std) was calculated.

| **[NaCl]** | **DF**  **(mAb2)** | ***〈R_h_〉_,app_* _replicate_1** | ***〈R_h_〉_,app_* _replicate_2** | **Average** | **Std** |
| --- | --- | --- | --- | --- | --- |
| **0** | 1 (45 mg/mL) | 4,6 | 4,6 | 4,6 | 0,00 |
| **0** | 2 | 4,7 | 4,6 | 4,7 | 0,07 |
| **0** | 4 | 4,7 | 4,8 | 4,8 | 0,07 |
| **0** | 8 | 4,9 | 4,8 | 4,9 | 0,07 |
| **0** | 16 | 4,9 | 4,8 | 4,9 | 0,07 |
| **0** | 32 | 5,0 | 4,9 | 5,0 | 0,07 |
| **10** | 1 (45 mg/mL) | 7,3 | 7,0 | 7,2 | 0,21 |
| **10** | 2 | 6,0 | 5,8 | 5,9 | 0,14 |
| **10** | 4 | 5,6 | 5,5 | 5,6 | 0,07 |
| **10** | 8 | 5,3 | 5,2 | 5,3 | 0,07 |
| **10** | 16 | 5,1 | 5,0 | 5,1 | 0,07 |
| **10** | 32 | 5,0 | 4,9 | 5,0 | 0,07 |
| **150** | 1 (45 mg/mL) | 6,2 | 6,1 | 6,2 | 0,07 |
| **150** | 2 | 5,4 | 5,5 | 5,5 | 0,07 |
| **150** | 4 | 5,3 | 5,2 | 5,3 | 0,07 |
| **150** | 8 | 5,1 | 5,1 | 5,1 | 0,00 |
| **150** | 16 | 5,0 | 5,0 | 5,0 | 0,00 |
| **150** | 32 | 4,9 | 4,9 | 4,9 | 0,00 |

**Table S3: Apparent hydrodynamic radii (*〈R_h_〉_,app_*) of a mAb2 dilution series (dilution factor (DF) 1 (corresponding to 45 mg/mL) to 32, as a function of NaCl concentration ([NaCl].** The average was determined from two independent samples (*〈R_h_〉_,app_* _replicate_1 and *〈R_h_〉_,app_* _replicate_2), from which the standard deviation (std) was calculated.

| **[NaCl]** | **DF** | ***〈R_h_〉_,app_* _replicate_1** | ***〈R_h_〉_,app_* _replicate_2** | **Average** | **Std** |
| --- | --- | --- | --- | --- | --- |
| **0** | 1 (45 mg/mL) | 2,3 | 2,2 | 2,3 | 0,07 |
| **0** | 2 | 2,4 | 2,4 | 2,4 | 0,00 |
| **0** | 4 | 3,0 | 3,0 | 3,0 | 0,00 |
| **0** | 8 | 3,7 | 3,7 | 3,7 | 0,00 |
| **0** | 16 | 4,2 | 4,2 | 4,2 | 0,00 |
| **0** | 32 | 4,6 | 4,6 | 4,6 | 0,00 |
| **10** | 1 (45 mg/mL) | 4,6 | 4,5 | 4,6 | 0,07 |
| **10** | 2 | 4,7 | 4,6 | 4,7 | 0,07 |
| **10** | 4 | 4,7 | 4,6 | 4,7 | 0,07 |
| **10** | 8 | 4,9 | 4,8 | 4,9 | 0,07 |
| **10** | 16 | 5,0 | 4,9 | 5,0 | 0,07 |
| **10** | 32 | 5,0 | 5,0 | 5,0 | 0,00 |
| **150** | 1 (45 mg/mL) | 6,6 | 6,5 | 6,6 | 0,07 |
| **150** | 2 | 5,9 | 5,7 | 5,8 | 0,14 |
| **150** | 4 | 5,5 | 5,4 | 5,5 | 0,07 |
| **150** | 8 | 5,3 | 5,2 | 5,3 | 0,07 |
| **150** | 16 | 5,2 | 5,1 | 5,2 | 0,07 |
| **150** | 32 | 5,1 | 5,1 | 5,1 | 0,00 |

**Table S4: Apparent hydrodynamic radii (*〈R_h_〉_,app_*) of a mAb3 dilution series (dilution factor (DF) 1 (corresponding to 45 mg/mL) to 32, as a function of NaCl concentration ([NaCl].** The average was determined from two independent samples (*〈R_h_〉_,app_* _replicate_1 and *〈R_h_〉_,app_* _replicate_2), from which the standard deviation (std) was calculated.

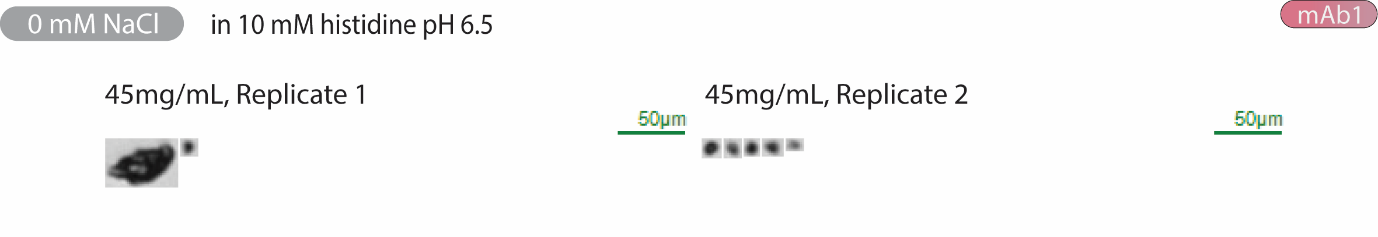

**Figure S1: MFI thumbnail collages of the sub-visible particles >5 µm detected in ~0.250 mL 45 mg/mL mAb1 in 10 mM histidine pH 6.5**

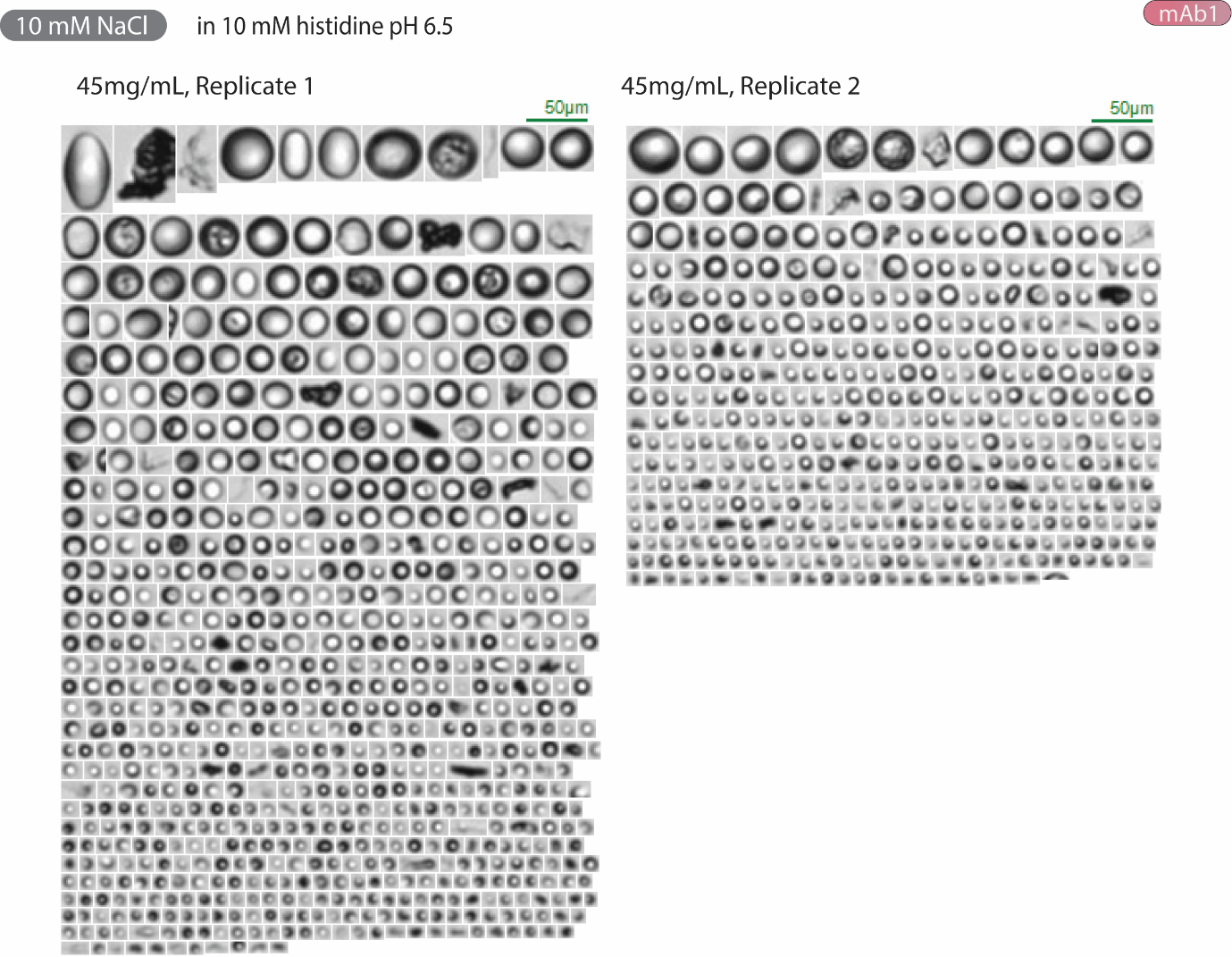

**Figure S2: MFI thumbnail collages of the sub-visible particles >5 µm detected in ~0.250 mL 45 mg/mL mAb1 + 10 mM NaCl in 10 mM histidine pH 6.5**

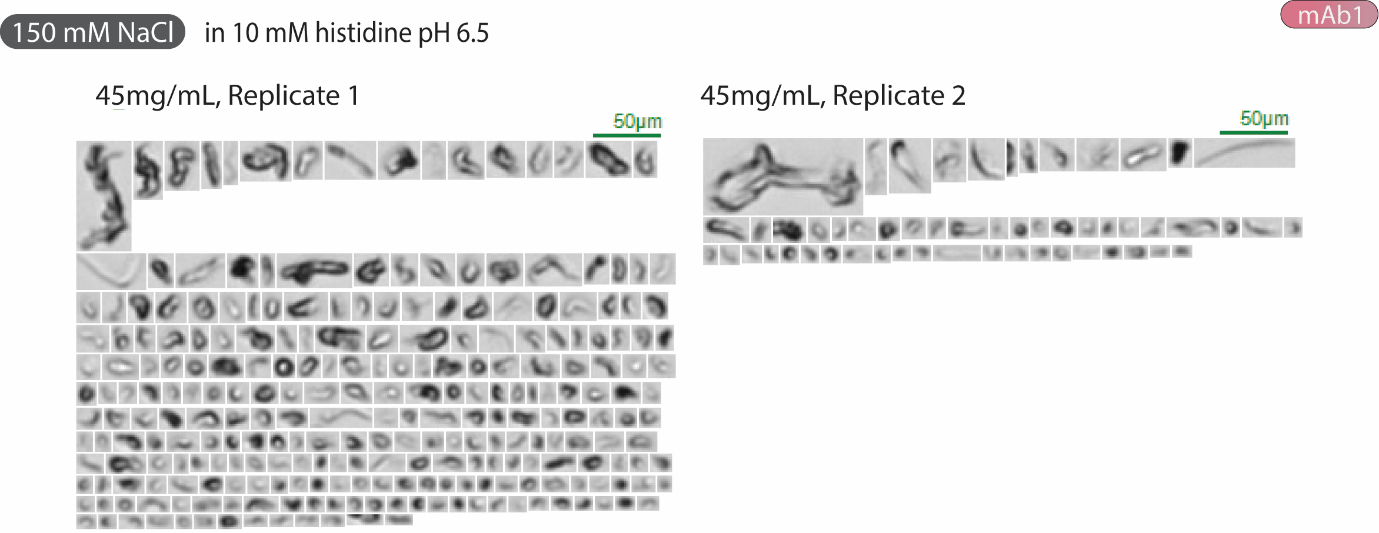

**Figure S3: MFI thumbnail collages of of the sub-visible particles >5 µm detected in ~0.250 mL 45 mg/mL mAb1 + 150 mM NaCl in 10 mM histidine pH 6.5**

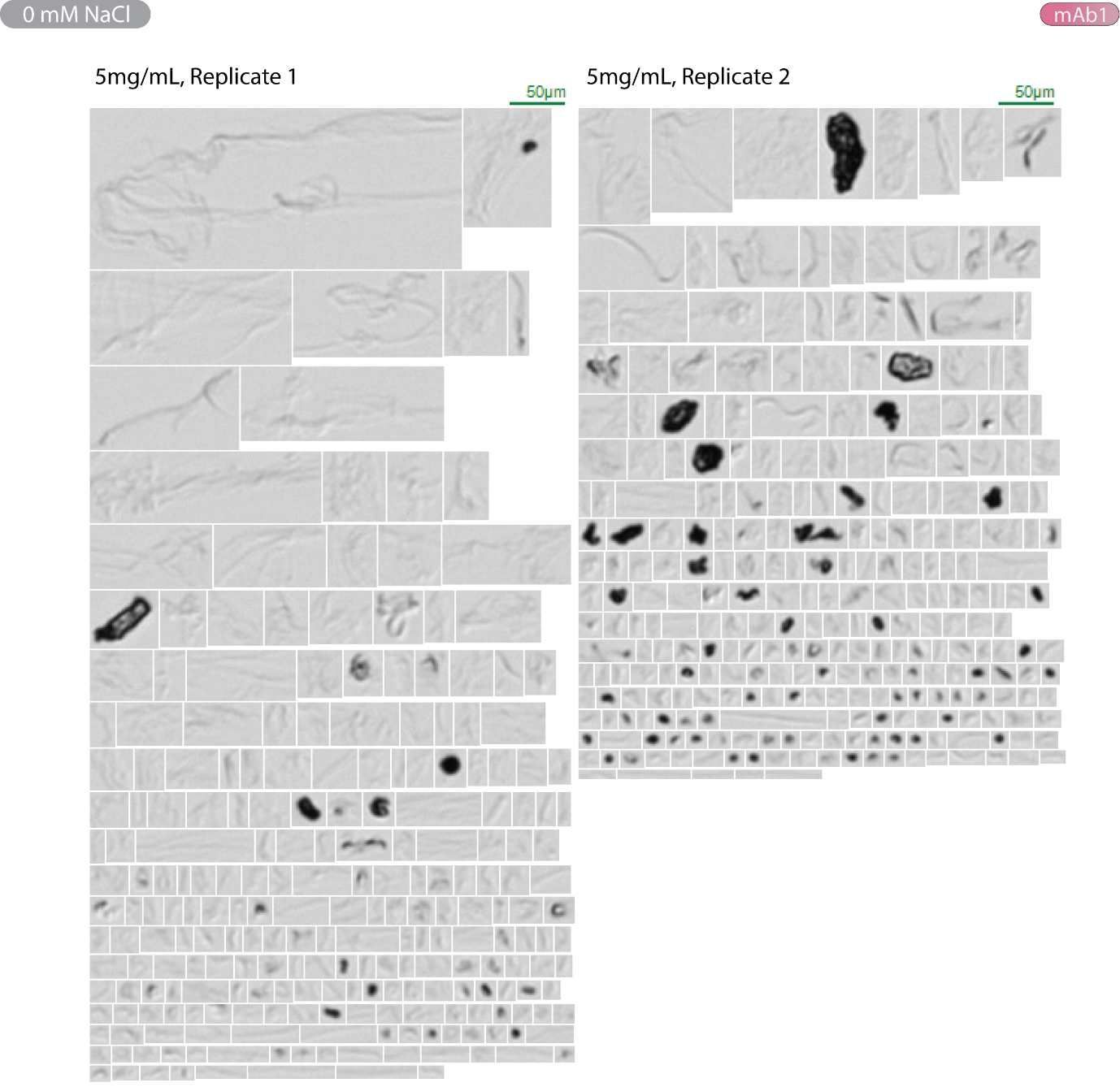

**Figure S4: MFI thumbnail collages of the sub-visible particles >5 µm detected in ~0.250 mL 5 mg/mL mAb1 in 10 mM histidine pH 6.5 (no salt)**

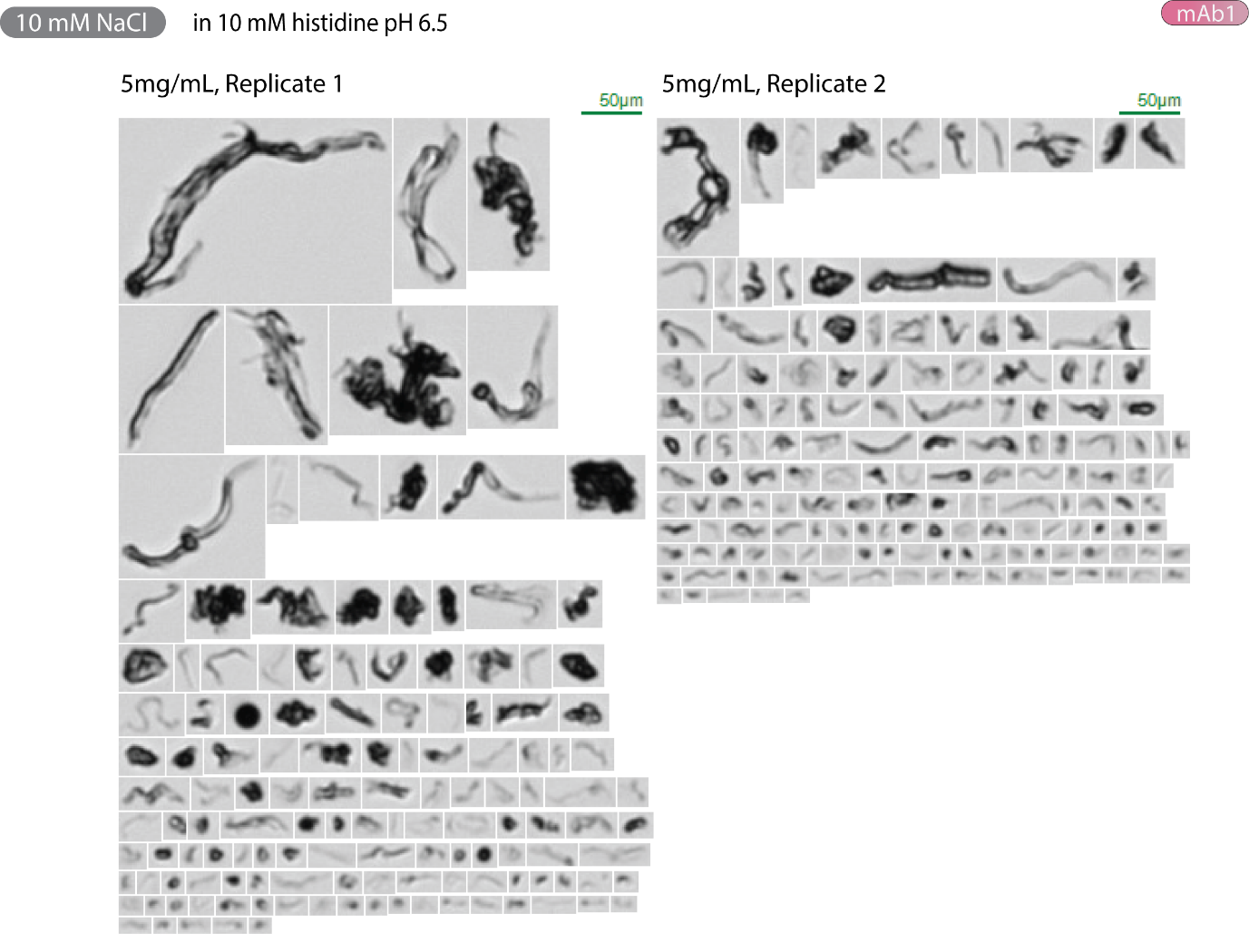

**Figure S5: MFI thumbnail collages of the sub-visible particles >5 µm detected in ~0.250 mL 5 mg/mL mAb1 +10 mM NaCl in 10 mM histidine pH 6.5**

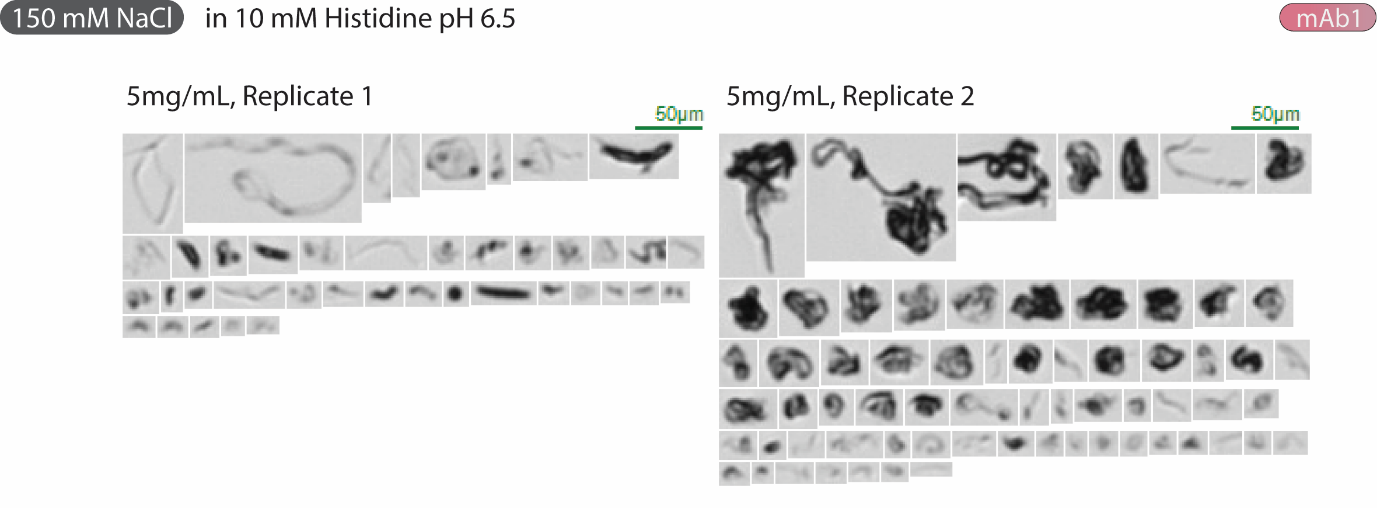

**Figure S6: MFI thumbnail collages of the sub-visible particles >5 µm detected in ~0.250 mL 5 mg/mL mAb1 +150 mM NaCl in 10 mM histidine pH 6.5**

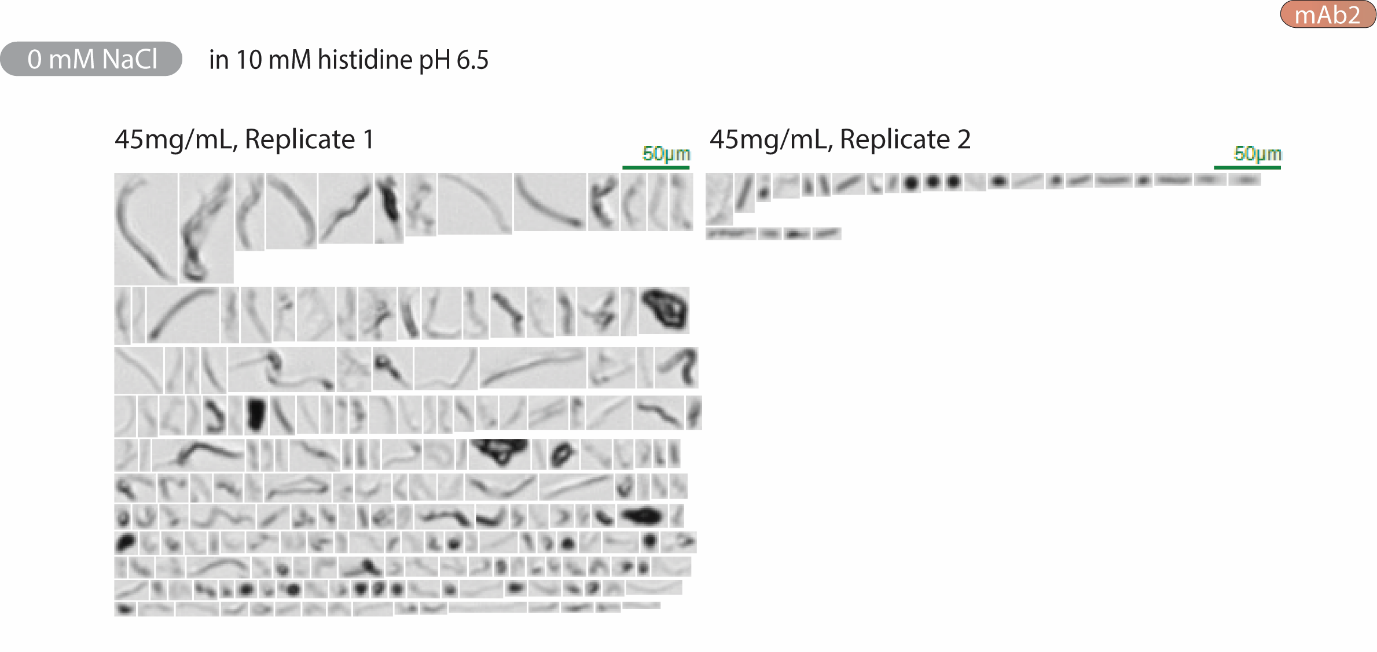

**Figure S7: MFI thumbnail collages of the sub-visible particles >5 µm detected in ~0.250 mL 45 mg/mL mAb2 in 10 mM histidine pH 6.5 (no salt)**

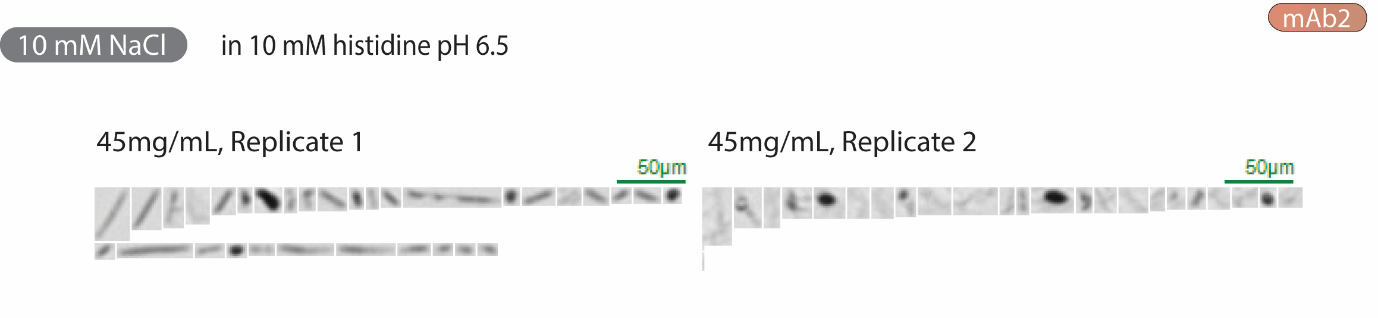

**Figure S8: MFI thumbnail collages of the sub-visible particles >5 µm detected in ~0.250 mL 45 mg/mL mAb2 + 10 mM NaCl in 10 mM histidine pH 6.5**

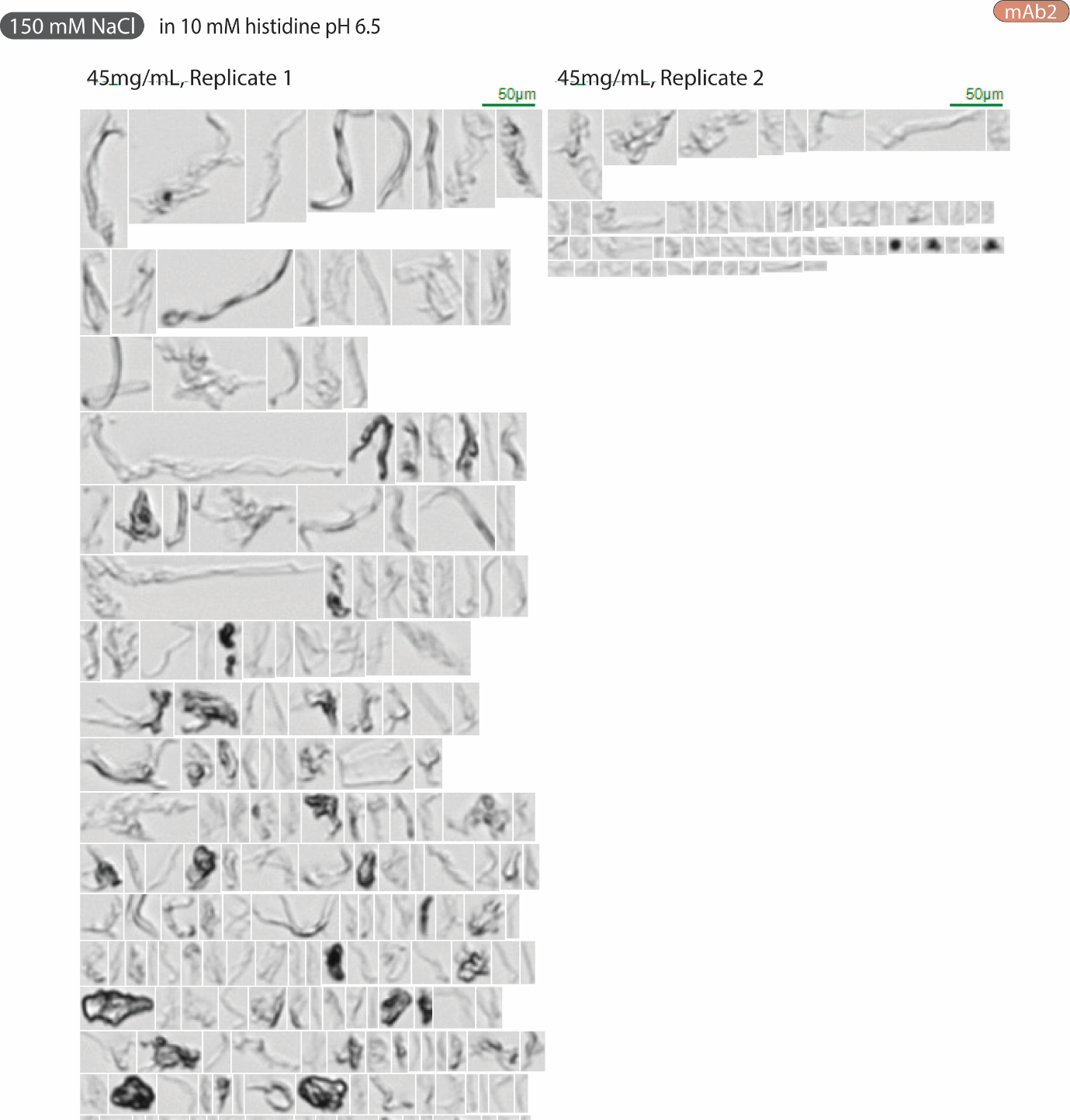

**Figure S9: MFI thumbnail collages of the sub-visible particles >5 µm detected in ~0.250 mL 45 mg/mL mAb2 + 150 mM NaCl in 10 mM histidine pH 6.5**

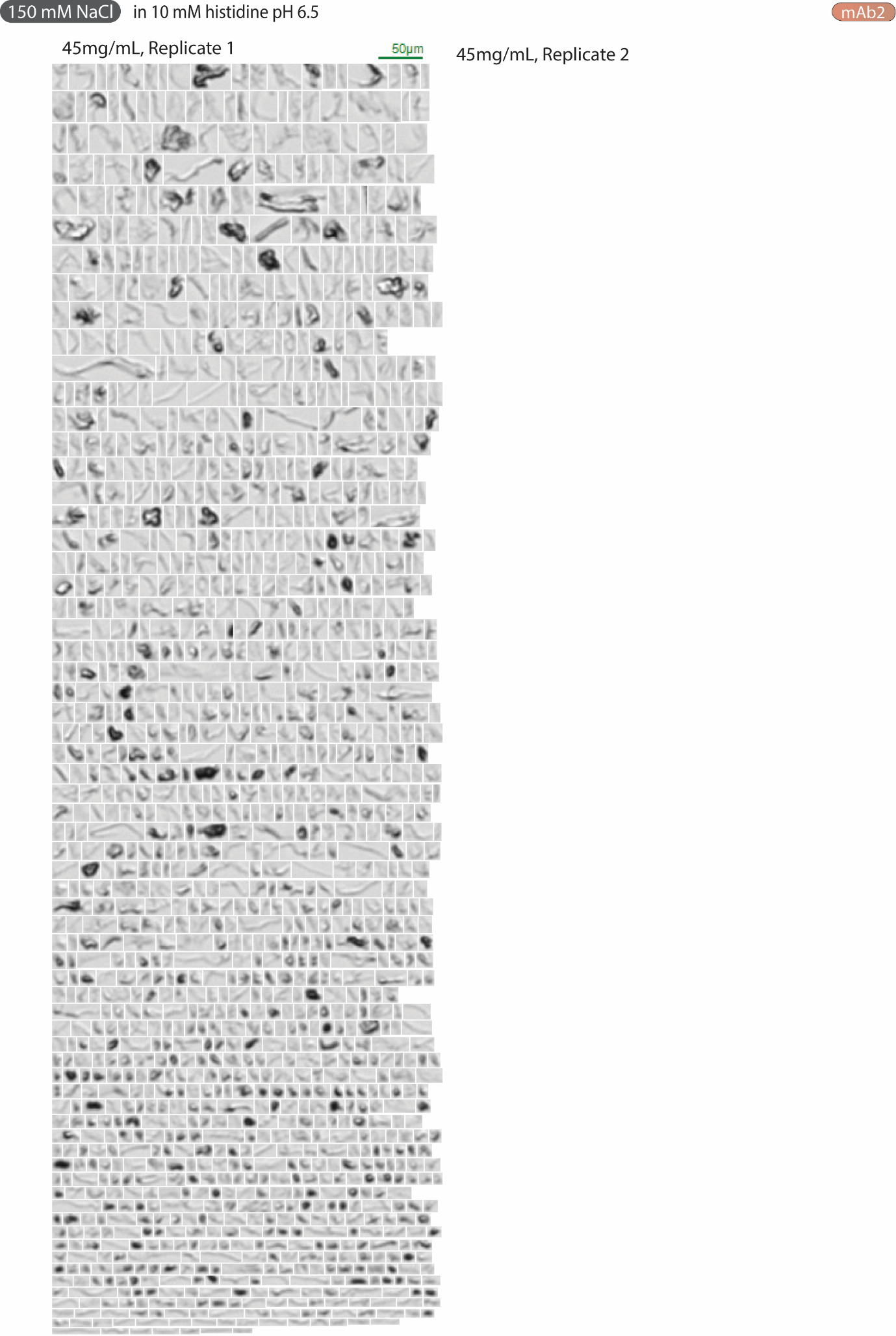

**Figure S10: MFI thumbnail collages of the sub-visible particles >5 µm detected in ~0.250 mL 45 mg/mL mAb2 + 150 mM NaCl in 10 mM histidine pH 6.5 (continued)**

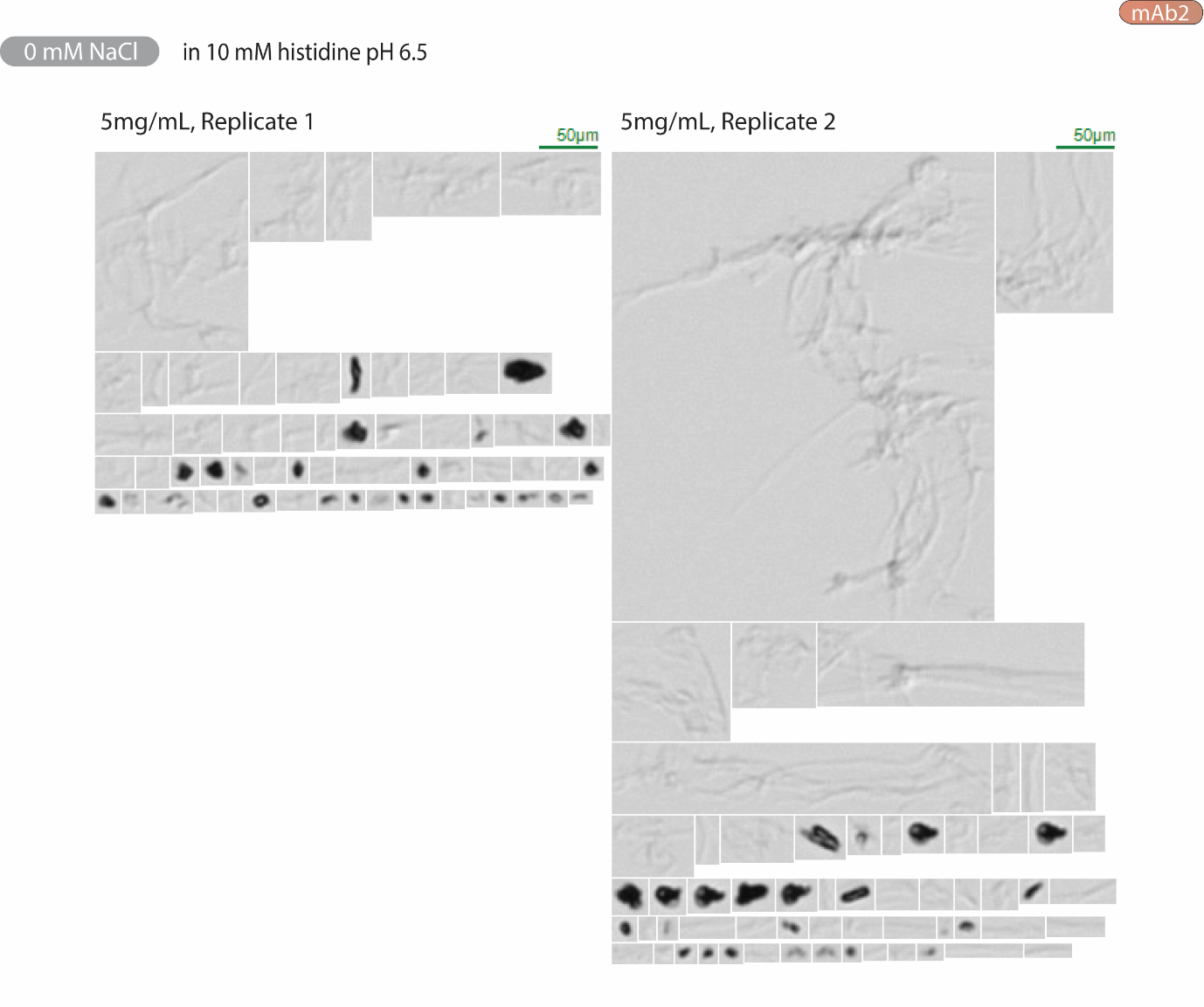

**Figure S11: MFI thumbnail collages of the sub-visible particles >5 µm detected in ~0.250 mL 5 mg/mL mAb2 in 10 mM histidine pH 6.5 (no salt)**

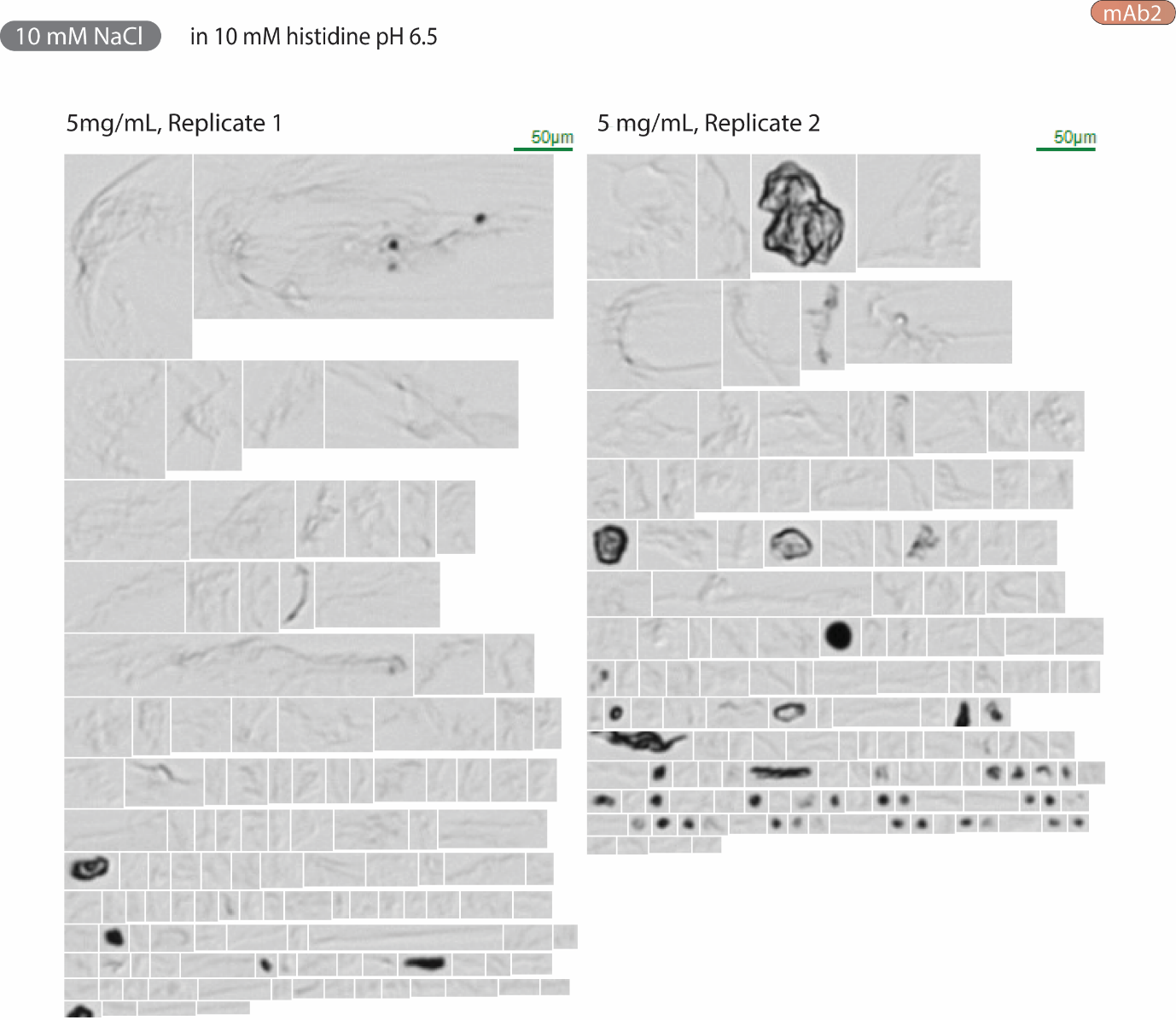

**Figure S12: MFI thumbnail collages of the sub-visible particles >5 µm detected in ~0.250 mL 5 mg/mL mAb2 +10 mM NaCl in 10 mM histidine pH 6.5**

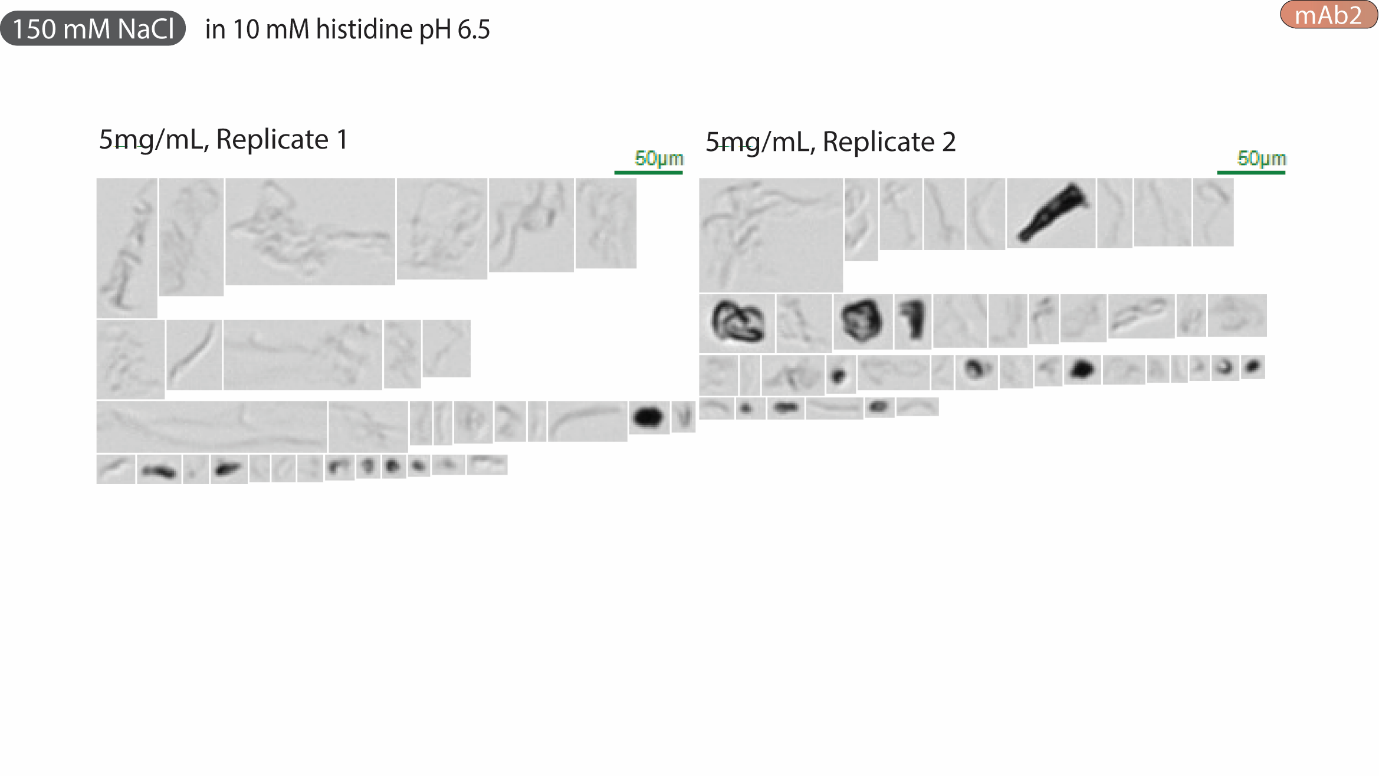

**Figure S13: MFI thumbnail collages of the sub-visible particles >5 µm detected in ~0.250 mL 5 mg/mL mAb2 +150 mM NaCl in 10 mM histidine pH 6.5**

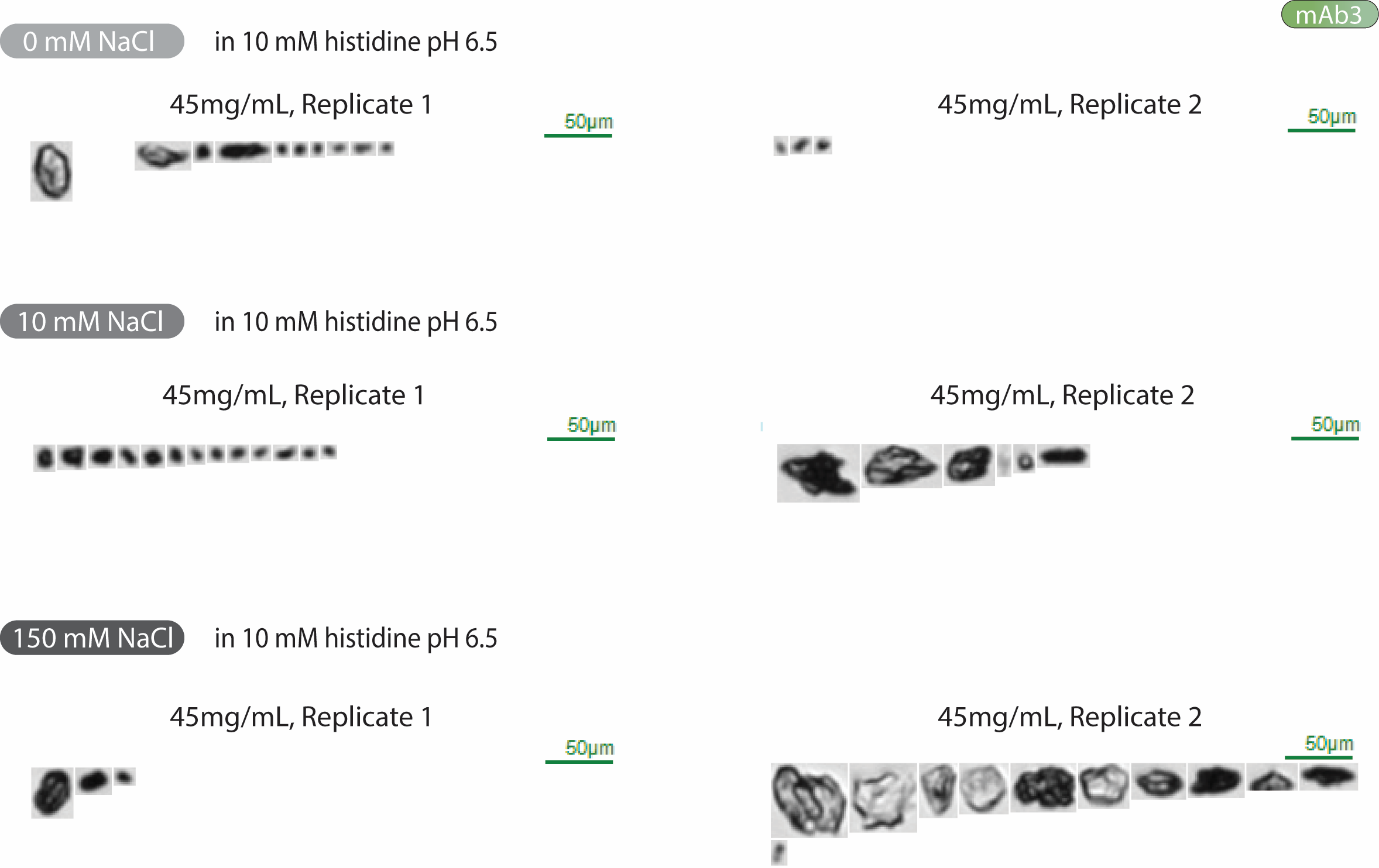

**Figure S14: MFI thumbnail collages of the sub-visible particles >5 µm detected in ~0.250 mL 45 mg/mL mAb3 +0/10150 mM NaCl in 10 mM histidine pH 6.5**

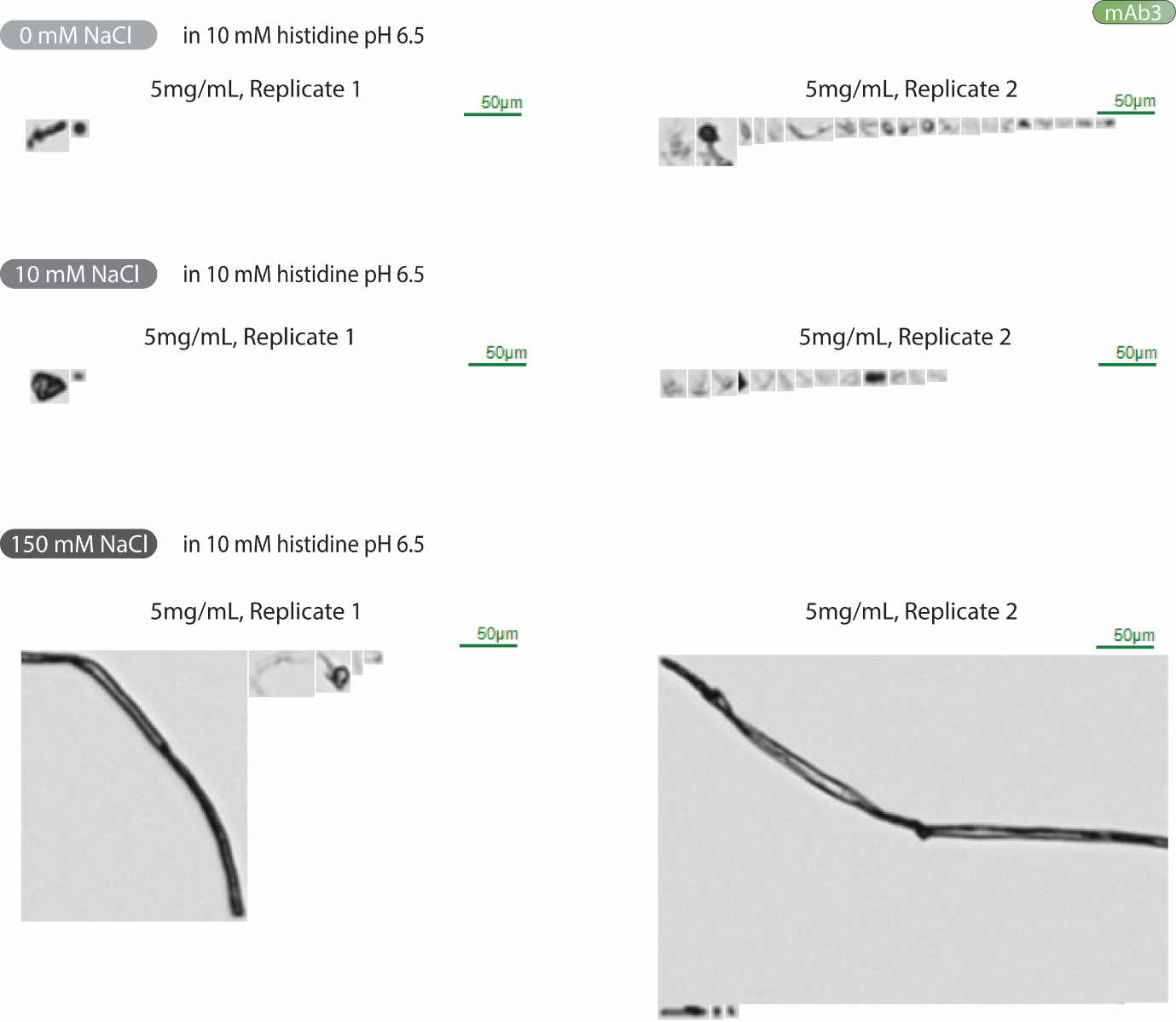

**Figure S15: MFI thumbnail collages of the sub-visible particles >5 µm detected in ~0.250 mL 5 mg/mL mAb3 +0/10/150 mM NaCl in 10 mM histidine pH 6.5**

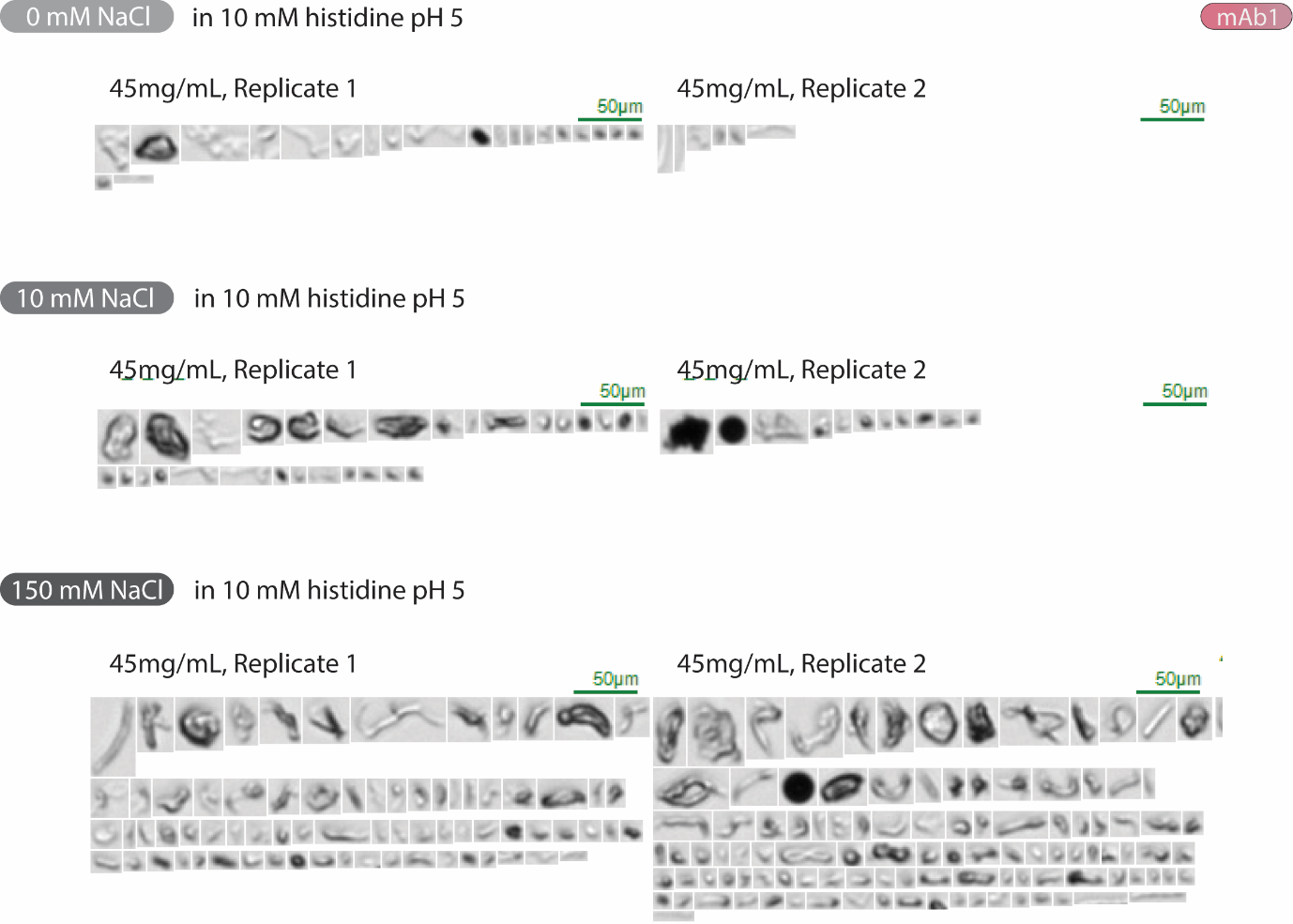

**Figure S16: MFI thumbnail collages of the sub-visible particles >5 µm detected in ~0.250 mL 45 mg/mL mAb3 +0/10/150 mM NaCl in 10 mM histidine pH 5**

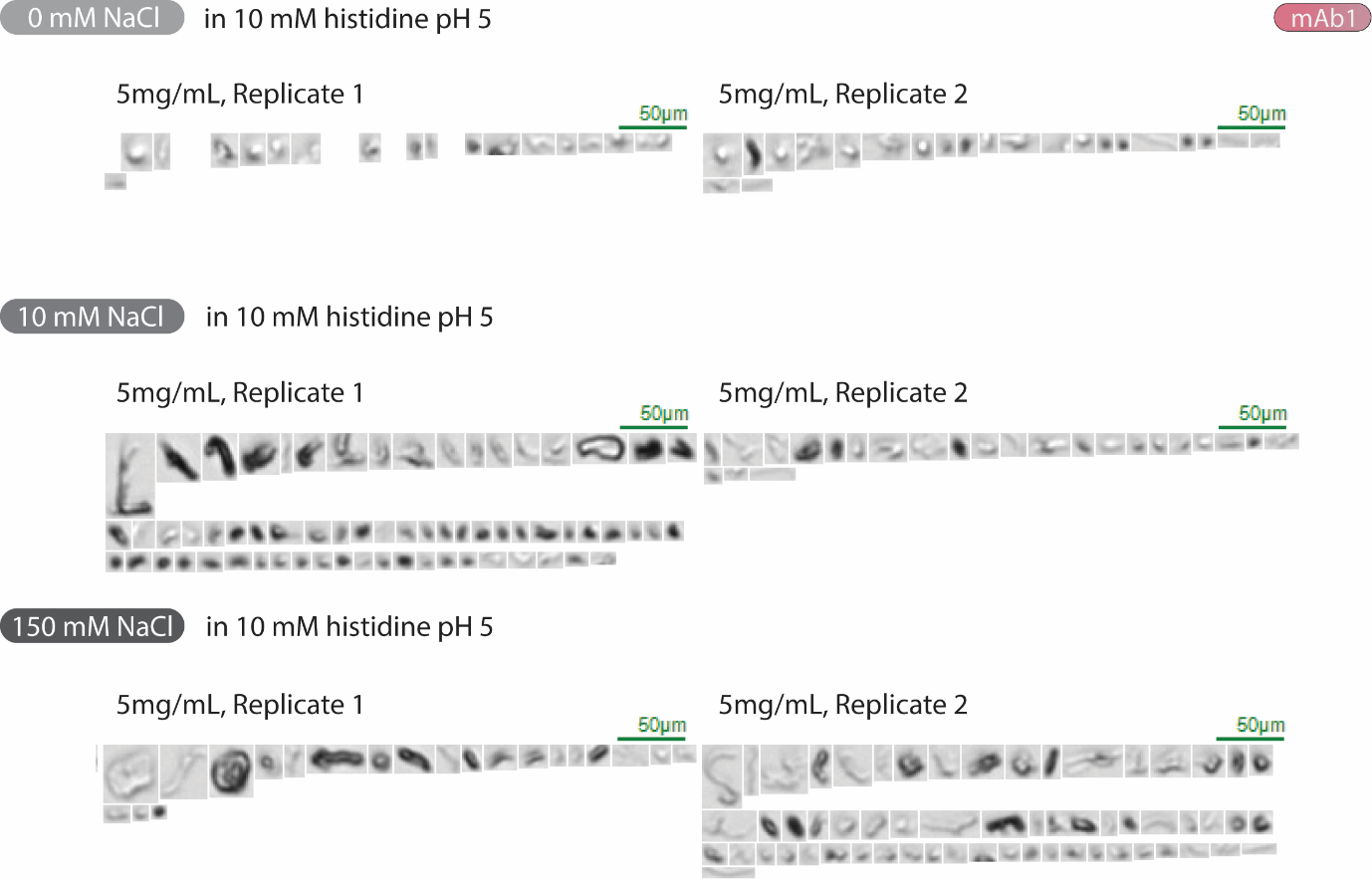

**Figure S17: MFI thumbnail collages of the sub-visible particles >5 µm detected in ~0.250 mL 5 mg/mL mAb3 +0/10/150 mM NaCl in 10 mM histidine pH 5**
